# Nepali oral microbiomes reflect a gradient of lifestyles from traditional to industrialized

**DOI:** 10.1101/2024.07.01.601557

**Authors:** Erica P. Ryu, Yoshina Gautam, Diana M. Proctor, Dinesh Bhandari, Sarmila Tandukar, Meera Gupta, Guru Prasad Gautam, David A. Relman, Ahmed A. Shibl, Jeevan Bahadur Sherchand, Aashish R. Jha, Emily R. Davenport

## Abstract

**Background:** Lifestyle plays an important role in shaping the gut microbiome. However, its contributions to the oral microbiome remains less clear, due to the confounding effects of geography and methodology in investigations of populations studied to date. Furthermore, while the oral microbiome seems to differ between foraging and industrialized populations, we lack insight into whether transitions to and away from agrarian lifestyles shape the oral microbiota. Given the growing interest in so-called ‘vanishing microbiomes’ potentially being a risk factor for increased disease prevalence in industrialized populations, it is important that we distinguish lifestyle from geography in the study of microbiomes across populations.

**Results:** Here, we investigate salivary microbiomes of 63 Nepali individuals representing a spectrum of lifestyles: foraging, subsistence farming (individuals that transitioned from foraging to farming within the last 50 years), agriculturalists (individuals that have transitioned to farming for at least 300 years), and industrialists (expatriates that immigrated to the United States within the last 20 years). We characterize the role of lifestyle in microbial diversity, identify microbes that differ between lifestyles, and pinpoint specific lifestyle factors that may be contributing to differences in the microbiomes across populations. Contrary to prevailing views, when geography is controlled for, oral microbiome alpha diversity does not differ significantly across lifestyles. Microbiome composition, however, follows the gradient of lifestyles from foraging through agrarianism to industrialism, supporting the notion that lifestyle indeed plays a role in the oral microbiome. Relative abundances of several individual taxa, including *Streptobacillus* and an unclassified Porphyromonadaceae genus, also mirror lifestyle. Finally, we identify specific lifestyle factors associated with microbiome composition across the gradient of lifestyles, including smoking and grain source.

**Conclusion:** Our findings demonstrate that by controlling for geography, we can isolate an important role for lifestyle in determining oral microbiome composition. In doing so, we highlight the potential contributions of several lifestyle factors, underlining the importance of carefully examining the oral microbiome across lifestyles to improve our understanding of global microbiomes.

## Introduction

Throughout the last 300,000 years, our species experienced continual cultural transformation marked by milestones such as the development and use of tools, specialized division of labor, and urbanization [1]. These cultural shifts profoundly influenced both human societies and biology. One major transition in recent human history was the shift in subsistence strategy from hunting and gathering to agriculture, and subsequently to industrialization. Such transitions encompass multifaceted lifestyle changes, including shifts in diet, population density, infectious disease burden, habitat, and other environmental factors [2]. These factors individually play pivotal roles in shaping the human microbiome — the diverse collection of bacteria, archaea, fungi and other eukaryotes, and viruses that inhabit our bodies [3–6].

Understanding the role of subsistence strategy and accompanying lifestyle transitions has become a major focus of microbiome research [7–10]. Numerous studies show that the gut microbiome shifts with industrialization. Specifically, industrialized populations generally exhibit lower gut microbiome alpha diversity compared to traditional populations, often lacking microbes commonly found in traditional populations, such as *Prevotella* and *Treponema* [5, 11–13]. These differences are associated with diet, drinking water source, and social structure [13–16]. While considerable progress has been made in understanding how the gut microbiome differs across lifestyles, differences in the oral microbiome across these transitions remain largely uncharacterized. Addressing this gap is crucial considering both the role of the oral microbiome in oral and systemic health [17–19], and also its prominence in the context of ancient DNA research [10, 20]. Therefore, it is important to expand studies of the oral microbiome to encompass diverse global populations practicing different subsistence strategies.

Much of our understanding of the oral microbiome across subsistence strategies comes from analyses of ancient dental calculus [20, 21]. Similar to bones, dental plaque can fossilize into calculus, preserving oral microbial communities for millennia [10, 22]. These ancient DNA samples have been utilized for investigating shifts in the oral microbiome over time, especially through major lifestyle transitions such as the Neolithic Revolution [23–26]. For example, the Neolithic Revolution was accompanied by increasing levels of putatively pathogenic microbes in the oral microbiome, which is believed to be due to increased starch consumption. While immensely valuable for directly observing historical microbiomes, the use of ancient DNA has its limitations. Specimens are extremely scarce, which often results in small sample sizes from a particular location and time period. Furthermore, given the rarity of samples, there are often considerable confounding temporal and geographical gaps between groups practicing different lifestyles. Finally, detailed information about specific lifestyle variables is often lacking beyond what can be inferred from the physical characteristics of the tooth or the archeological artifacts recovered from the site of collection.

The oral microbiomes of modern humans living a range of lifestyles offer an alternative approach. While much of our understanding of the oral microbiome stems from industrialized populations, predominantly of European descent [27, 28], a limited number of studies have explored the oral microbiomes of non-industrialized populations [29–34]. These studies report that oral microbial diversity decreases with industrialization, with microbiome composition exhibiting differences based on lifestyle as well. More specifically, relative abundances of *Neisseria*, *Haemophilus, Prevotella,* and *Streptococcus* tend to decrease with industrialization [30, 31, 33, 35, 36]. However, several potential confounding factors may underlie the reported differences. First, geography is associated with the microbiome [13, 37–39]. Distinguishing between the potential effects of geography versus lifestyle remains challenging, given the often large geographic variability between populations practicing different lifestyles. Second, some studies incorporate publicly available microbiome data from industrialized individuals with newly generated sequencing data from traditional populations [29, 31]. Technical discrepancies in sample collection, processing, and sequencing can influence microbiome study outcomes, raising questions about whether observed variations are due to lifestyle or technical factors [40, 41]. Lastly, studies that effectively control for technical effects are primarily focused on traditional populations. As a result, we lack an understanding of how the oral microbiome differs across the entire spectrum of lifestyles, from traditional to industrialized, while adequately controlling for temporal, geographical, and technical variation.

To address these gaps, we characterized the salivary microbiota of Nepali individuals across a spectrum of human subsistence strategies, from traditional foragers to agriculturalists. The Chepang represent foragers, the Raji and Raute are hunters and gatherers that recently settled and began subsistence farming in the 1980s, the Tharu and Newars living in Nepal are agriculturalists, and the Nepali expatriates living in the United States represent a population that recently transitioned to industrialization. We also include Americans of European descent as representatives of a fully industrialized population. We demonstrate that oral microbiome composition differs along a gradient of traditional to industrialized lifestyles but, unlike that of the gut microbiome, differences are relatively subtle. By integrating questionnaire-based data encompassing diverse lifestyle variables such as diet, education, medical practices, we identify specific lifestyle factors associated with oral microbial compositional changes. Finally, we examine the gut-oral microbiome axis to evaluate whether the degree of intra-individual similarity between the two sites differs across lifestyles. These results demonstrate that like the gut microbiome, the oral microbiome mirrors lifestyle.

## Results

### Description of Populations

We investigated the oral microbiome in diverse Nepali populations practicing a spectrum of lifestyles and an American population representing industrialized lifestyle. Briefly, the Nepali individuals in this study belonged to five ethnic groups native to Nepal - Chepang (n = 18), Raji (n = 11), Raute (n = 14), Tharu (n = 20), and Newar (n = 8) (**Fig 1, S1 Table**). We also included expatriate Newar (n = 12) and European-Americans (n = 6), both of whom reside in the San Francisco Bay Area. The Chepang, numbering 84,400 [42], are foragers that primarily reside in small communities of remote, isolated villages within the hills of the lower Himalaya in central Nepal. The Chepang village in this study lacks modern amenities such as electricity, running water, and other indicators of urbanization. While they supplement their diets with food grown via slash and burn agriculture, low productivity of the hilly terrain compels them to heavily depend on foraged jungle foods such as undomesticated tubers and wild nettle (*sisnu*). The Raji and Raute, previously nomadic foragers, transitioned to subsistence agriculture within the past 50 years. The Raute and Raji are among the smallest ethnic groups of Nepal. The Raute, with roughly 550 individuals [42], reside in the far western hills of Nepal, whereas the Raji, numbering 5,100 [42], inhabit the neighboring Terai plains. These two populations will be referred to as “recently settled” in this study, due to the transitional nature of their lifestyle. In contrast, the Tharu and Newar, two of the largest ethnic groups in Nepal numbering 1.8 and 1.3 Million respectively [42], practice agriculture. The Tharu, hailing from the Terai plains in Southern Nepal, fully transitioned to agriculture about 300 years ago. The Newar originate from Kathmandu valley and are renowned for their cultural and economic contributions to Nepal. Although increasing urbanization of Kathmandu valley has afforded some Newar individuals access to industrialized comforts, the population in this study resides in a relatively rural village in the outskirts of Kathmandu valley and primarily engages in agriculture. For those reasons, in this study, both the Tharu and Newar will be referred to as the established agriculturalists. The expatriate Newar in the US (“expats”) also originated from the Kathmandu valley and emigrated within the past 20 years, settling in the United States in their mid-30s. The Chepang, Raji, Raute, and Tharu individuals sampled here largely overlap with a previous study of Nepali gut microbiomes [15]. Both the fecal samples in this previous study and saliva samples in this current study were collected concurrently. By focusing on individuals across a range of lifestyles within a confined geographic region, our study aims to discern oral microbial signatures of lifestyle without the confounding effects of geography, climate, and technical factors.

**Figure 1:**
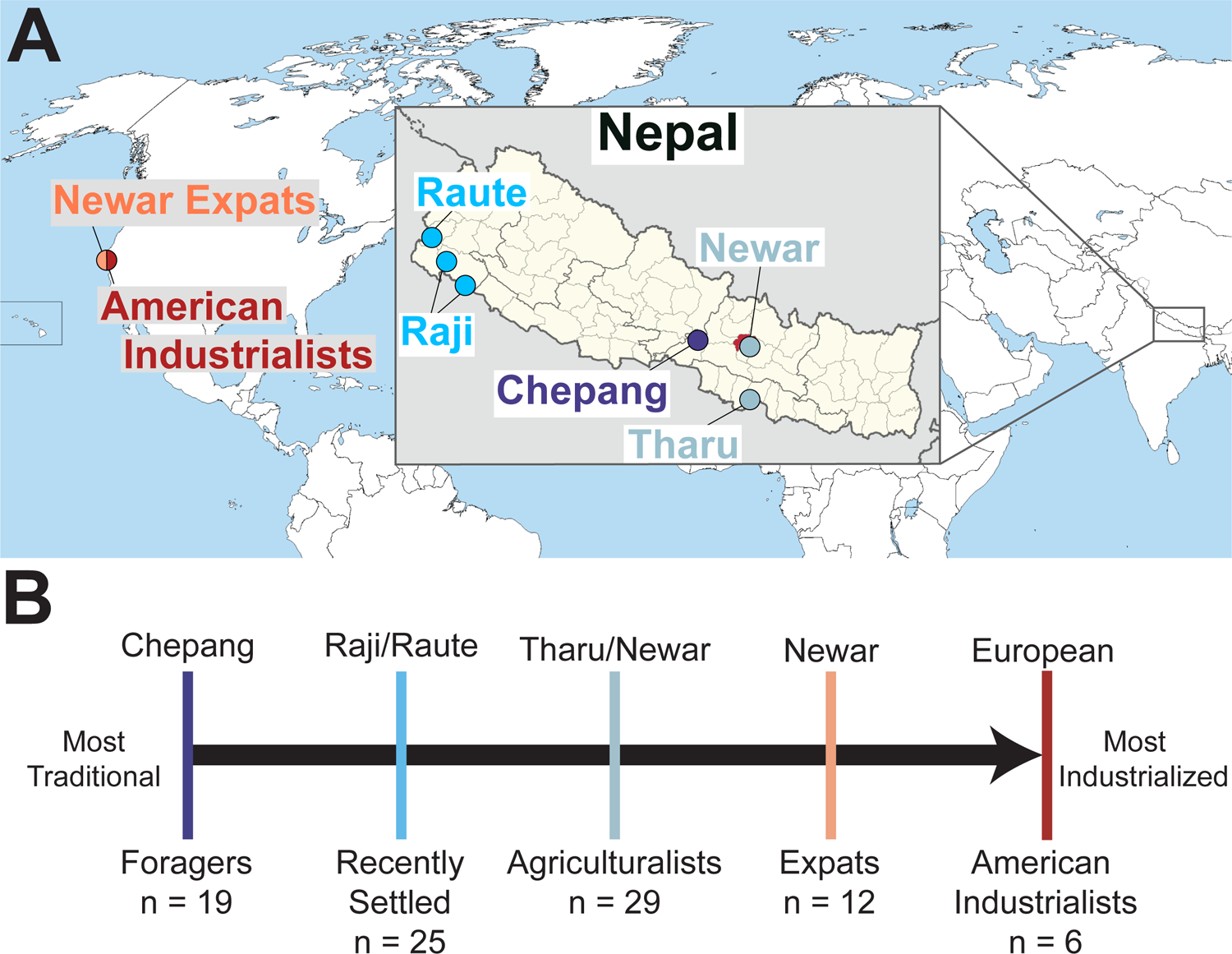
Sampling locations of all Nepal- and US-based Populations A) Locations of the populations sampled in Nepal and the US. The US populations are specifically from the Northern California region. Location of Kathmandu is indicated in red on the Nepal map. Colors correspond to lifestyle groupings as described in (B). B) Oral microbiome samples were collected from individuals that span a spectrum of lifestyles, from nomadic foraging populations (dark blue), to populations that recently transitioned from foraging to small scale agriculture (teal), to established small scale agriculturalists (sky blue), to Nepali expats residing in the US practicing an industrialized lifestyle (peach), to American Industrialists (red). Sample sizes for each lifestyle category are indicated.

### Lifestyle factors differ across populations

Given that multiple dietary, environmental, and socio-economic factors differ across populations in Nepal, we administered a survey to capture the specific lifestyle factors that differ across the Nepali populations (**S1 Table**). We used Random Forests to evaluate the ability of the 37 lifestyle factors obtained from the survey questions to classify individuals to their respective lifestyle groups. Nepali individuals are correctly assigned to their respective lifestyle categories with 92.06% overall accuracy (**S1 Fig**), suggesting that the populations in our study are highly distinguishable based on these factors.

We then performed correspondence analysis (CA) to determine which specific variables distinguish the lifestyles. Our analysis reveals significant dependence between the samples and lifestyle factors (p = 0.004; Chi-square test of independence). Individuals cluster closely in the top two dimensions of CA based on their lifestyle group, as do similar specific lifestyle variables (ie. education and literacy, **S2 FigA**). Specifically, CA axis 1 follows the lifestyle trend (p = 3.26×10^-14^; Jonckheere-Terpstra test), whereas CA axis 2 does not (p = 0.21; Jonckheere-Terpstra test, **S2 FigB-C**). The top 10 contributing lifestyle factors to CA1 include *sisnu* consumption, fuel source, and literacy (**S2 FigD**). For example, the Chepang consume *sisnu* most frequently, use solid fuel for cooking, and have low literacy rates, while the Expats do not consume *sisnu*, use gas or electricity for cooking, and are literate. In contrast, the top contributing factors to CA2 include behavioral factors like smoking and alcohol consumption (**S2 FigE**).

### Oral microbiome diversity does not differ across Nepali populations

Fecal samples concurrently collected with the saliva from the Chepang, Raji, Raute, and Tharu individuals previously revealed pronounced gut microbiome compositional differences across the gradient of lifestyles from foraging to industrialized [15]. To evaluate whether oral microbiome compositional differences align with the continuum of lifestyles, we initially characterized the oral microbiome via saliva samples collected from 89 individuals across 5 lifestyles (**S1 Table**). Recognizing the potential for DNA extraction methodology to introduce variability in microbiome studies [43, 44], we adopted multiple approaches to ensure robustness of our conclusions. For each sample, DNA extraction was performed using two different kits: Qiagen QIAamp MinElute Virus Spin kit and MO BIO PowerSoil DNA. These kits were selected as the MO BIO Powersoil DNA extraction kit was commonly used in microbiome studies, while the Qiagen QIAamp MinElute Virus Spin kit was recommended by DNA Genotek for oral microbiome DNA collection. Both extracts were sequenced in the same sequencing run at comparable sequencing depths. Overall microbiome composition and diversity are consistent between the two kits (**S3 Fig**). Consequently, all subsequent analyses were performed using the data obtained from the Qiagen kit for simplicity, which resulted in 69 individuals that passed quality control steps (see Methods).

We first evaluated whether overall microbiome diversity differed across lifestyle groups, as decreasing diversity is typically thought of as a hallmark of traditional to industrialized lifestyle transitions [12, 31, 33]. We observe no significant difference in Shannon diversity across the lifestyle groups (p > 0.05, Kruskal-Wallis; **Fig 2**), but there is a significant difference in Faith’s Phylogenetic diversity between the lifestyles (p = 0.028; Kruskal-Wallis, **Fig 2**). A post-hoc pairwise comparison demonstrates that the American Industrialists are driving the differences in Faith’s Phylogenetic diversity (American Industrialists vs. other lifestyles: p < 0.05, Dunn’s post-hoc test). Notably, there is no significant difference in Faith’s Phylogenetic diversity between the four Nepali populations (p > 0.05; Kruskal-Wallis). These findings remain largely consistent with other alpha diversity metrics across both extraction kits (see Methods, **S4 Fig, S5 Fig**). Aligning with the observations in the gut microbiomes of these individuals [15], our results indicate that oral microbiome diversity does not correlate with lifestyle differences within Nepal when geography is controlled for. Notably, our sample sizes, while modest, are larger than most other oral microbiome studies examining traditional lifestyles [29–31, 33], underscoring that the lack of signal is not due to insufficient power.

**Figure 2:**
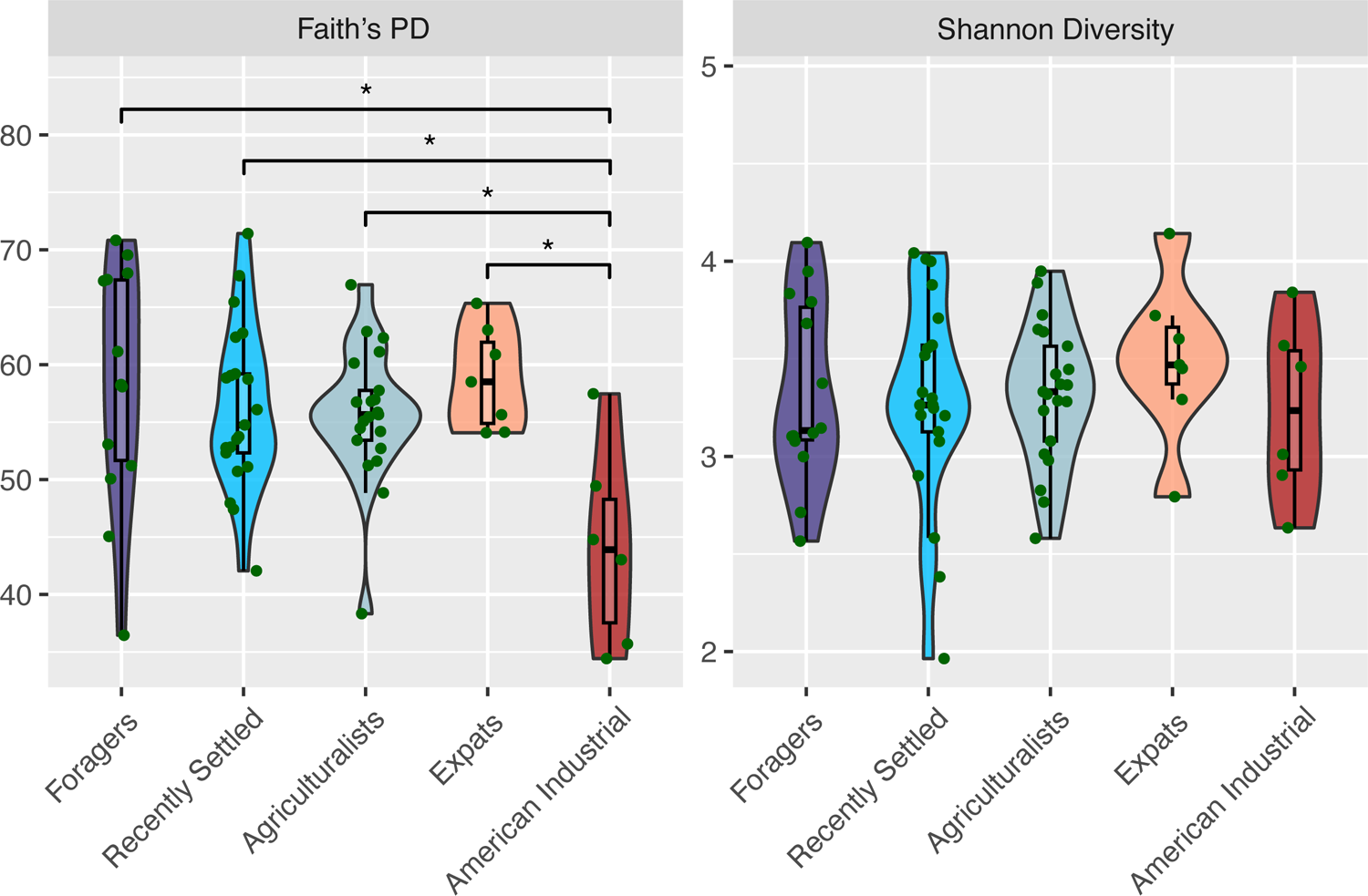
Alpha diversity does not significantly differ by lifestyle in Nepal. Faith’s phylogenetic diversity (Faith’s PD - left) and Shannon alpha diversity (Shannon Diversity - right) shown for all individuals, grouped by lifestyle. Lifestyles are ordered from most traditional (Foragers) to most industrialized (American Industrialists), left to right. No significant difference detected across lifestyles for Shannon alpha diversity (p = 0.8, Kruskal-Wallis), but a marginally significant difference detected for Faith’s phylogenetic diversity (p = 0.028, Kruskal-Wallis). Notably, those significant differences only occur between the American Industrialists and other lifestyle groups, not between Nepali individuals residing in Nepal or the US. Significant differences (p < 0.05) between specific populations are indicated (*).

### Oral microbiome composition differs across lifestyles

Unlike microbiome diversity, microbiome composition varies across lifestyles. We calculated between-sample Bray-Curtis distances to measure beta diversity (Bray & Curtis, 1957), revealing that oral microbiome composition varies significantly with lifestyle (p= 2.3×10^-4^; PERMANOVA, **Fig 3A**). When visualized via Principal Coordinate Analysis (PCoA), the first axis (PCoA1, explaining 28.62% of microbiome variation) follows the lifestyle gradient (p = 0.0014; Jonckheere-Terpstra test, **Fig 3B**), with the Expatriates and Americans being most different from the traditional Nepali populations. This pattern is consistent across data from both extraction kits (**S6 FigB-D**) and with UniFrac metrics (**S7 Fig, S8 Fig**) [46]. These compositional differences across groups, however, are fairly subtle, with classification of microbiomes into lifestyle groupings via Random Forests not being better than expected by random chance (43.48% accuracy, **S9 Fig**).

**Figure 3:**
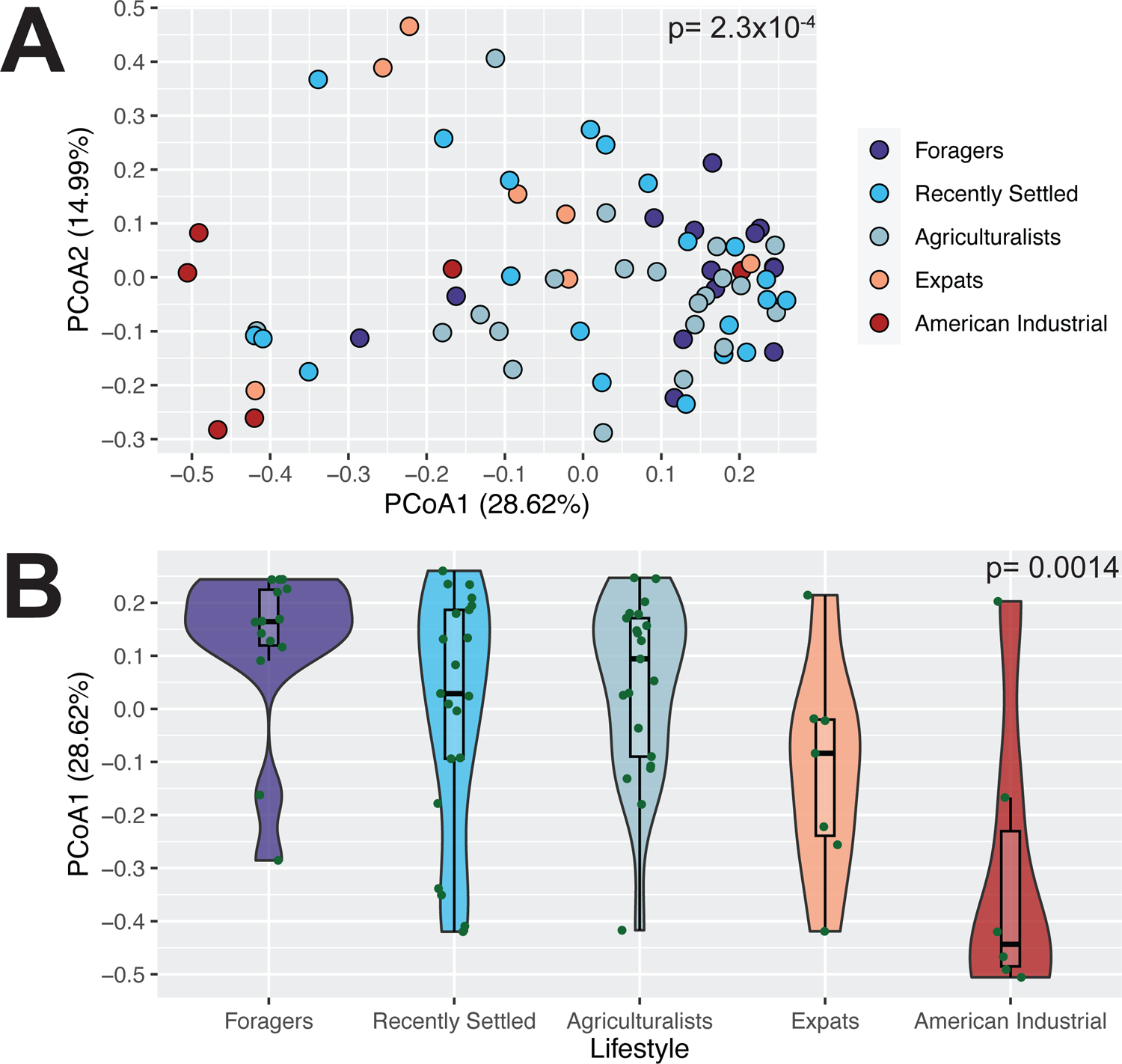
Oral microbiome composition significantly differs based on lifestyle. A) Microbiome composition varies significantly with lifestyle (p = 2.3×10^-4^, PERMANOVA). The PCoA plot shows individuals ordinated based on Bray-Curtis distance and colored by lifestyle. B) The distribution of individuals along PCoA axis 1 follows the lifestyle gradient, from traditional to Industrial (p = 0.0014, Jonckheere-Terpstra test). Lifestyles are ordered from most traditional (Foragers) to most industrialized (American Industrialists), left to right.

Because overall microbiome composition differed based on lifestyle, we next determined which specific taxa differed across the lifestyles. To do this, we conducted differential abundance analysis using ALDEx2, which accounts for compositionality in its application [47]. We observe that 2 of the 111 oral genera (1.8%) were significantly differentially abundant after accounting for multiple tests, namely *Streptobacillus* and an unclassified Porphyromonadaceae genus (padj = 0.011 and 0.021, respectively; Kruskal-Wallis) (**S2 Table**). We also implemented an alternative approach for identifying which taxa were following the lifestyle gradient by performing the Jonckheere-Terpstra test for all genera and correcting for multiple tests. We find that nine genera significantly followed the lifestyle gradient, including the two that were also identified using ALDEx2 (**Fig 4, S3 Table**). Eight genera - *Streptobacillus*, *Porphyromonadaceae_unclassified, Granulicatella, Moraxella, Simonsiella, Neisseria, Bacteroidetes_unclassified, and Brachymonas -* show decreasing abundance with industrialization, consistent with the trend observed in Faith’s Phylogenetic diversity. The only exception is *Atopobium*, which shows the opposite trend of increasing abundance with industrialization (**Fig 4**).

**Figure 4:**
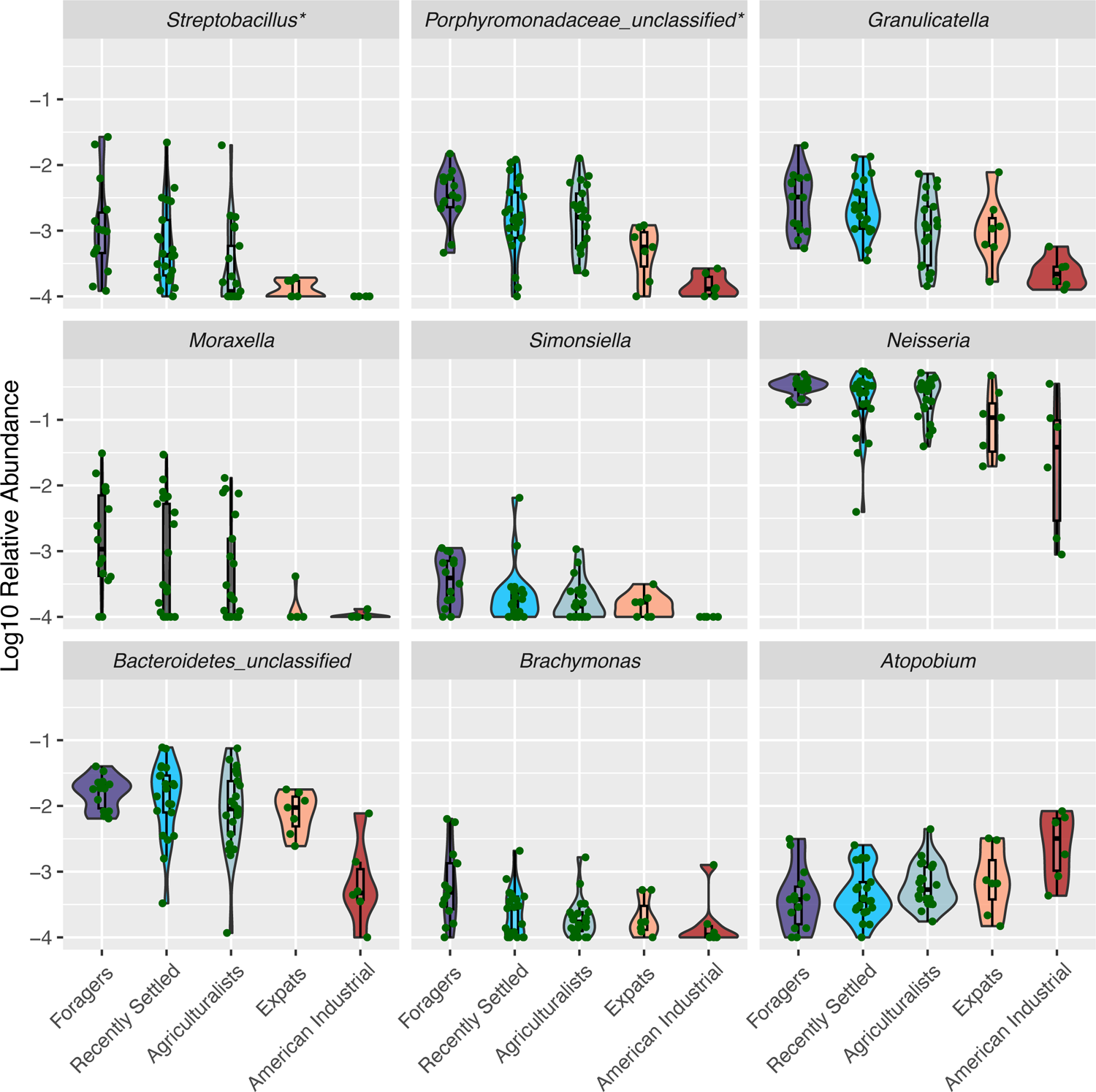
Abundances of nine genera significantly follow the lifestyle gradient. The relative abundances of nine genera significantly follow the lifestyle gradient via a Jonckheere-Terpstra test followed by Benjamini-Hochberg correction (adjusted p < 0.05). Lifestyles are ordered from most traditional (Foragers) to most industrialized (American Industrialists), left to right. All taxa have been log10-transformed for visualization purposes. Taxa marked with * are also significantly differentially abundant across lifestyles based on ALDEx2. Most taxa tend to decrease in relative abundance as the lifestyles transition from more traditional to industrial, while the abundance of *Atopobium* increases.

### Grain type is associated with microbiome differences across lifestyles

Given that overall microbiome composition mirrored the transition of lifestyles within Nepal, we sought to identify lifestyle factors that potentially underlie these differences. To do this, we first started at a broad scale by comparing the major axes defining oral microbiome composition and the lifestyle variables, from PCoA and CA respectively. We calculated the correlation between the first three CA axes, which cumulatively captured 37.14% of variation in the lifestyle survey data, and first three PCoA axes, which cumulatively captured 49.35% of variation in the microbiome data. We observe a significant correlation between PCoA2 and CA2, which is primarily comprised of behavioral lifestyle factors like tobacco and alcohol use and distinguishes the recently settled populations from foragers and agriculturalists (p = 0.03; rho = −0.27; Spearman correlation, **S10 FigA**). No significant correlation is observed between the CA axes and alpha diversity (**S10 FigB**).

We then identified which specific lifestyle factors are associated with the observed differences in microbiome composition. As testing all 37 measured lifestyle factors would be prohibitive due to multicollinearity, we selected the top 15 key lifestyle distinguishing factors based on their contributions to the first two CA axes (**S2 Fig**). We used these factors to perform canonical correspondence analysis (CCA) to determine which variables are associated with shifts in the microbiome. We find significant associations between these lifestyle distinguishing factors and the oral microbiome composition among the Nepalis (p = 0.013; ANOVA); with the top factors being alcohol consumption, smoking habits, location, *sisnu* consumption, and grain type (p = 0.044, 0.001, 0.003, 0.003, 0.027; respectively, ANOVA) (**Fig 5A**).

**Figure 5:**
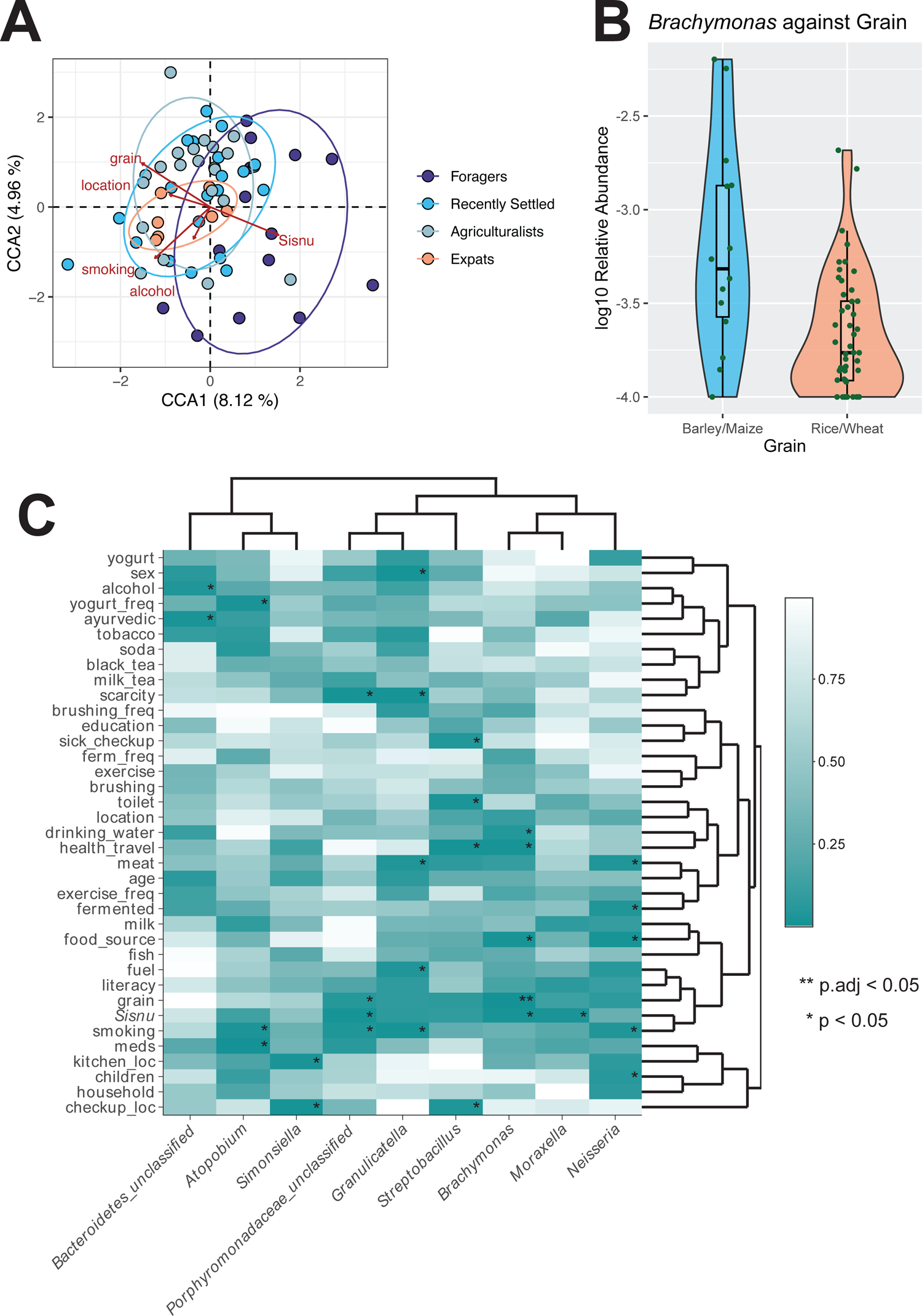
Alcohol, smoking, location, *sisnu*, and grain type are associated with the oral microbiome A) There are significant associations between lifestyle factors and the microbiome as observed via CCA. Points represent individuals and color represents corresponding lifestyle, with ellipses around each population. Red arrows represent the lifestyle factors significantly associated with the microbiome. A total of 15 lifestyle factors were inputted into the CCA model based on contribution to each CA axis. B) *Brachymonas* is significantly associated with grain type consumed (padj = 0.0219). Specifically, relative abundance is higher in individuals who report barley and maize consumption compared to rice and wheat. Taxa were log10 transformed for visualization. C) Several specific lifestyle factors are associated with individual oral genera. Significant associations based on a nominal p-value threshold are indicated with *. The significant association based on an adjusted p-value threshold is indicated with **.

Finally, we identified associations between the nine taxa that are differentially abundant across lifestyle groupings (**S3 Table**) and the 37 lifestyle factors included in our survey via linear models. Out of the 333 associations tested, we find that *Brachymonas* is significantly associated with grain consumption at an adjusted p-value, with higher abundance of this taxon observed in individuals who primarily consume barley and maize compared to those who primarily consume rice and wheat (padj = 0.022; **Fig 5B)**. Furthermore, we observe an additional 27 associations that are significant at a nominal p-value < 0.05 (**Fig 5C, S4 Table**). For example, we observe that the relative abundances of *Granulicatella*, *Neisseria,* and *Porphyromonadaceae_unclassified* are higher in non-smokers, whereas *Atopobium* relative abundance is lower in non-smokers (p = 0.006, p = 0.032, p = 0.033, p = 0.023; respectively, **S11 FigA**). Similarly, the relative abundances of *Brachymonas, Moraxella*, and *Porphyromonadaceae_unclassified* are higher in individuals that consume *sisnu* (p = 0.014, p = 0.019, p = 0.012; respectively, **S11 FigB**). Overall, these results demonstrate that a variety of lifestyle factors potentially underlie the differences in oral microbiome composition observed between lifestyles within Nepal.

### Predicted metabolism pathways are differentially abundant across lifestyles

In addition to the taxonomic differences observed between lifestyle groups, predicted functional potential of the microbiome significantly differs as well. Based on the use of PICRUSt2 [48], predicted functional abundance significantly varies with lifestyle (p = 0.0036; PERMANOVA, **S12 FigA)**. The top two PCA axes both significantly follow the lifestyle gradient (PC1: p = 0.049, PC2: p = 0.0064; Jonckheere-Terpstra test, **S12 FigB**). We note that while the taxa present in Nepali oral microbiomes seem to be fairly closely represented in the reference taxonomy used to predict gene content (average NSTI = 0.037), there are many caveats to predictive methods such as PICRUSt and these results should be viewed as hypothesis generating [49].

To identify specific potential functional differences across lifestyles, we conducted differential abundance testing with ALDEx2 using the predicted abundances of 109 pathways. Although none are significant after multiple test corrections, 22 pathways are significant at a nominal p-value of p < 0.05 (**S5 Table**), 13 of which are classified as metabolism pathways (**S13 FigA**). These metabolism pathways can be categorized into 7 classes, some of which increase in abundance with increasing industrialization, including lipid metabolism and glycan biosynthesis, while others decrease, such as xenobiotics degradation and microbial metabolism in diverse environments (**S13 FigB**). General transporter proteins (ATP-binding cassette transporters - padj = 0.0013, phosphotransferase system - padj = 0.0013) and degradation pathways (aminobenzoate degradation - padj = 0.0028) are significantly enriched via enrichment analysis (**S14 Fig**). Finally, the top 10 most significant pathways from ALDEx2 were further examined to identify the top contributing microbes and whether they differ by lifestyle. *Fusobacterium* is one of the top taxa contributing to platinum resistance (**S15 FigA**). There is significant enrichment of *Fusobacterium* in the traditional Nepali populations compared to industrialized populations (p = 0.0037; Kruskal-Wallis test, **S15 FigB**). Overall, predicted metabolism pathways significantly differ across lifestyle, mirroring the taxonomic gradient across lifestyles in Nepal.

### Microbial network structure varies across lifestyles

We then investigated network structure to determine whether community structure differs across the lifestyles. We used the SparCC module in the SpiecEasi package [50, 51] to generate a network from all 111 genera observed in this study. The resulting network consists of 37 nodes with at least one edge and 6 co-abundance groups, with a modularity of 0.45 (**Fig 6A**). Among the taxa identified as following the lifestyle gradient, 5 out of the 9 are connected to at least one other taxon, with *Porphyromonadaceae_unclassified, Neisseria, Bacteroidetes_unclassified*, and *Granulicatella* being in the same co-abundance group (CAG1), whereas *Atopobium* is in a separate co-abundance group (CAG2). Interestingly, the proportions of CAGs differ across lifestyles, with CAG1 decreasing with industrialization and CAG2 increasing with industrialization (**Fig 6B**). These results demonstrate that community network structure differs along the lifestyle gradient, in addition to individual microbial taxa and predicted functional potential.

**Figure 6:**
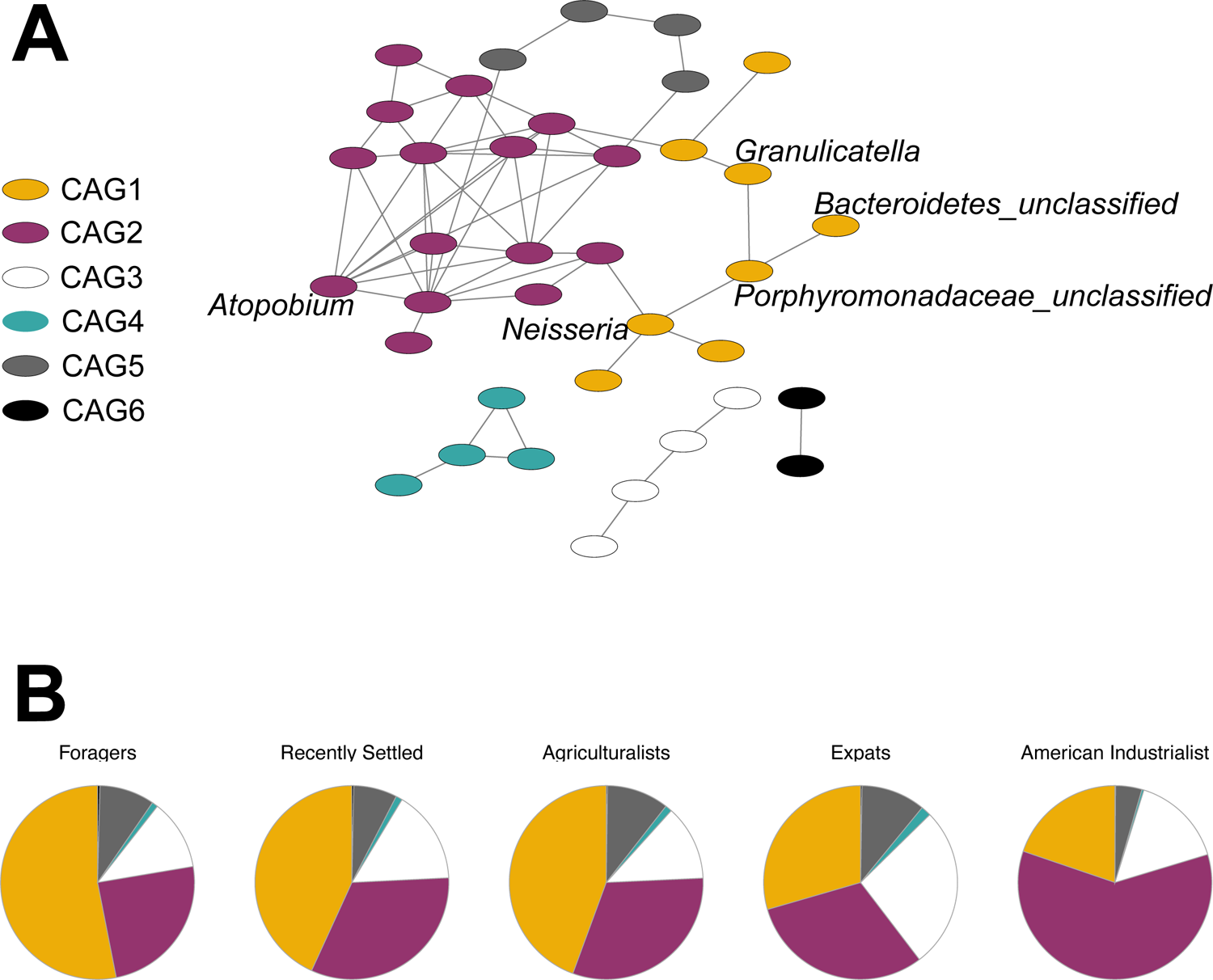
Differentially abundant taxa are highly connected in the oral microbiome co-occurrence network A) The SparCC module in the SpiecEasi package was used to generate a network from 111 genera. Network of 37 nodes with at least one significant edge is shown, with 6 co-abundance groups (CAGs) indicated by node color. Labeled nodes indicate genera that were identified as significantly differentially abundant across lifestyles. B) Proportions of CAGs vary across lifestyle. Specifically, CAG1 decreases with industrialization, whereas CAG2 increases with industrialization.

### Oral-gut microbiome distance decreases with agrarianism

Finally, we examined the role of lifestyle along the oral-gut microbiome axis. While different lifestyle factors independently associate with the oral and gut microbiomes, compositional similarity between the two sites within an individual increases with the extent of urbanization [32]. Thus, we were interested in assessing whether there was a similar association across lifestyles, in which we might expect to see increasing intra-individual similarity across the gradient of lifestyles from traditional to agrarian. To do this, we examined individuals for whom both oral and gut microbiome data were collected concurrently, for a total of 12 Foragers, 14 Recently Settled individuals, and 12 Agriculturalists (**S6 Table**), and calculated Bray-Curtis dissimilarity between the oral and gut microbiomes. We find that intra-individual oral-gut microbiome dissimilarity decreases, and therefore similarity increases, across the gradient of traditional to agrarian lifestyle as predicted, although this trend is not statistically significant (p = 0.11; Jonckheere-Terpstra test, **Fig 7**). We, however, do observe significant similarities in composition between the two body sites across individuals (p = 0.013, rho = −0.4; Spearman correlation, **S16 Fig**), suggesting that we are perhaps underpowered to detect significance of intra-individual dissimilarity at our current sample size.

**Figure 7:**
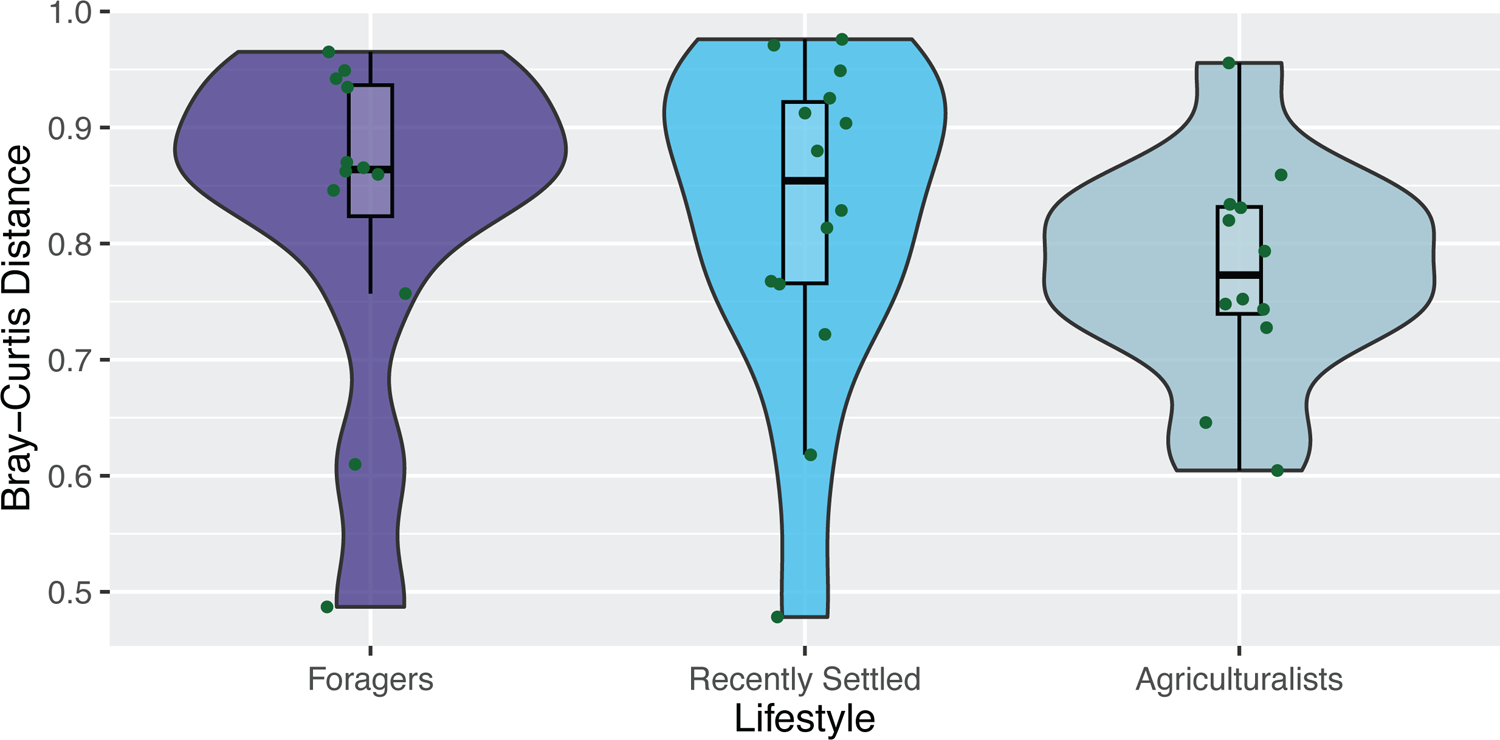
Correlation between the oral and gut microbiomes within an individual strengthens with agrarianism Microbiome dissimilarity (as measured by Bray Curtis dissimilarity) between the gut and oral microbiomes within an individual decreases across the gradient of lifestyles from traditional to agrarian for individuals residing in Nepal, albeit not significantly (p = 0.11; Jonckheere-Terpstra test).

## Discussion

Much of human microbiome research is weighted towards populations living in North America and Europe, with South Asia being particularly underrepresented [27]. As a consequence, our understanding of how the oral microbiome varies across human lifestyles is extremely limited, especially in regards to non-industrialized lifestyles including foraging, hunting and gathering, and small-scale agriculturalism. Some existing studies identify differences in the oral microbiome between hunter-gatherers and farmers [31, 33]. Others investigate the role of a few specific lifestyle factors; like smoking, alcohol consumption, and diet; but only in a single, usually industrialized, population [52–55]. Our study is the first comprehensive examination of the oral microbiome across multiple lifestyles, including transitional lifestyles – recently settled nomadic foraging individuals now practicing small scale agriculture and expatriates transitioning from farming to industrialization – and numerous lifestyle factors characterizing each lifestyle while controlling for geography and technical factors. We observe that even when controlling for geography, microbiome composition mirrors lifestyle, from foraging to industrialists, likely due to differences in dietary and behavioral habits.

In general, populations in industrialized countries have lower gut microbiome diversity than those practicing more traditional lifestyles [5, 11–14, 16, 56, 57]. This has led to an active discussion regarding whether we should intervene to improve human health and reduce microbial loss in industrialized countries [9, 58, 59]. While it is tempting to attribute the observed diversity differences across populations to lifestyle, controlling for geography eliminates those differences [14, 60, 61]. In fact, when examining the gut microbiomes of the individuals included in this study, there was no significant difference in within-sample alpha diversity across lifestyles, even when including American Industrialists [15]. Fewer studies examine oral microbiome diversity across human lifestyles, and most of them observe a decrease in diversity across lifestyles [31, 33]. That said, it is important to highlight that geography was not closely controlled for and might have confounded the results, as examining lifestyle while controlling for geography results in no difference in diversity [29].

Here, we demonstrate that alpha diversity of the oral microbiome does not significantly differ across lifestyles between Nepali individuals, aligning with findings from oral microbiome studies that control for geography [29]. Interestingly, unlike the gut microbiomes of the same individuals [15], we observe a significant decrease in alpha diversity in the American Industrial population compared to Nepali individuals, including the Nepali expats. As the Nepali expats and American Industrialists currently reside within the same metro area, geography is effectively controlled for. There are several possible explanations for why these differences are observed between the Nepali expats and American Industrialists, but not the populations within Nepal. First, the main differentiating factor may not be geography, as one might assume if only comparing the populations residing in Nepal to American Industrialists, but rather lifestyle factors that have more extreme effects between Industrialists and other lifestyles than between the traditional and agrarian populations within Nepal. For example, the Nepali expats included in this study tend to retain their traditional cuisine, which differs greatly from a standard American diet. Although recipes are modified to account for local ingredient availability, the main dietary components remain consistent across the ethnically Nepali populations, regardless of geography. Another possibility is that geography does drive oral microbial diversity, but can only do so during critical windows earlier in life in which oral microbiomes are malleable [62]. While gut microbiota appear to be malleable even with immigration well into adulthood [63], it is not clear if the same is true for the oral microbiome. Once a stable microbial community or host immune repertoire is established, moving to another geographic region may not result in diversity changes. The Nepali expatriates included in this study immigrated around their mid-30s, so they may have missed this window. Further investigation would be needed to tease apart these and other possible explanations for the differences in diversity we observed between Nepali Individuals and American Industrialists.

Similar to the gut microbiomes in the same individuals [15], oral microbiome composition mirrors lifestyle. Specifically, we observe a consistent compositional gradient when comparing individuals from traditional foraging populations (Chepang), to recently settled populations (Raji and Raute), to small-scale agriculturalists (Newar and Tharu), to immigrants (Newar) and industrialized Americans. Compared to the gut, which we reanalyzed using ALDEx2 to ensure comparability of results (**S7 Table**), differences across lifestyles are more muted in the oral microbiome. More genera in the gut are significantly differentially abundant across lifestyles (27% in gut, 1.8% in oral), even though there was slightly more power with the oral samples due to increased sample size. This finding may be due to increased resiliency of the salivary microbiome compared to the gut microbiome, thus resulting in fewer differences in the microbiome across lifestyle [64–68]. It is important to note that these considerations are specific to the salivary microbiome, as the microbiomes of other oral anatomical sites, including dental calculus, differ significantly from that of saliva [69–71].

When examining microbial abundance across lifestyle, most of the differentially abundant microbes decrease across the gradient of traditional to industrialized. Many of these taxa co-occur and lie in the same co-abundance group when considering the full oral microbiome network (CAG1). One such taxon is *Neisseria,* which decreases in abundance with industrialization in other lifestyle studies [31, 35]*. Neisseria* plays a beneficial role in periodontal health, possibly by preventing the colonization of pathogenic microbes [72, 73]. Its decreasing abundance aligns with the hypothesis that the loss of crucial microbes is associated with the emergence of disease in industrialized society [74]. Other previously identified lifestyle-associated oral microbes are not significantly associated with lifestyle in our study, such as *Haemophilus, Prevotella,* and *Streptococcus* [30, 31, 33, 36]. Instead, we observe decreasing levels of an unclassified Porphyromonadaceae genus. *Porphyromonas gingivalis* is a member of the Porphyromonadaceae family, well-established as a pathogenic oral microbe contributing to periodontal disease [75]. Further investigation is needed of this microbe in our dataset, as this would provide insight into *Porphyromonas* as a potential oral pathogen at a global scale. *Streptobacillus* is another microbe that decreases in abundance with increasing industrialization. In North America, *Streptobacillus* is most well-known as *Streptobacillus moniliformis* for its presence in rat oral microbiomes and role in rat bite fever [76]. In contrast, *Streptobacillus* appears as a commensal member in the human oral microbiomes of Asian populations. More specifically, *Streptobacillus hongkongensis* resides in the oral cavity of populations from Hong Kong and the United Arab Emirates [77, 78]. *Streptobacillus* has also been found in the Agta hunter-gatherers in the Philippines [79], thus suggesting that *Streptobacillus* presence in the human oral microbiome may be regionally limited to Asia. This hypothesis aligns with our study, as *Streptobacillus* is not observed in the American Industrialists. The relative abundances of differentially abundant microbes decreasing with industrialization is consistent with prior observations of decreased microbiome diversity in industrialized populations.

In contrast, the one genus that significantly increases in relative abundance with industrialization is *Atopobium.* High levels of oral *Atopobium* relative abundance are associated with a variety of negative health outcomes, many of which are more common in industrialized societies. Highly abundant in individuals with dental caries, *Atopobium* is believed to contribute to development of dental caries as an accessory to *Streptococcus mutans*, a leading microbial cause of caries [80–82]. Oral *Atopobium* carriage is also enriched in individuals with hypertension [83], Sjögren’s Syndrome [84], and patients with severe oral mucositis, a toxicity occurring from cancer treatments [85]. Notably, we observe that *Atopobium* belongs to a separate co-abundance group from the other differentially abundant taxa, CAG2. CAG2 also contains *Veillonella,* a microbe that may be an accessory for oral pathogen colonization [86]. Future research is needed to disentangle whether *Atopobium* and other potentially pathogenic microbes play causal roles in the development of oral conditions.

Multiple specific lifestyle factors are believed to play a role in shaping the oral microbiome, such as smoking, dietary fiber, and carbohydrate source [53, 54, 87]. Most of these factors, however, were characterized in industrialized populations and it remains unclear whether the same factors play a role in the oral microbiota of traditional populations. Using extensive survey metadata, we observe significant associations between the microbiome and 15 lifestyle factors, with smoking, grain type, and *sisnu* consumption most strongly associated.

Smoking consistently associates with oral microbiome composition across numerous industrialized populations, with increases in *Atopobium* and decreases in *Neisseria* and family level Porphyromonadaceae being trademark indicators [54, 88–91]. We identify similar associations with smoking in our study as well. Specifically, *Atopobium* abundance is increased in smokers, whereas *Granulicatella*, *Neisseria,* and *Porphyromonadaceae_unclassified* abundances are decreased in smokers. The taxa with increased abundances in smokers cluster in the same co-abundance group, whereas *Atopobium* is found in a separate co-abundance group. These findings suggest that smoking habits may play an important, consistent role in defining oral microbiome community dynamics across lifestyles and highlight the importance of accounting for smoking as a factor in future studies of lifestyle and the oral microbiome.

The associations we observe between grain type and the oral microbiome in Nepali populations is of particular interest, given the proposed importance of starch-rich foods in shaping oral microbiomes along the primate lineage [92]. Carbohydrates are associated with a myriad of oral microbes, either due to their role in starch digestion or pathogenicity [93, 94]. In this study specifically, populations reported primarily consuming either barley and maize, or rice and wheat. Barley and maize differ substantially from rice and wheat in terms of phenol content, digestibility, fiber content, and glycemic index [95, 96]. More specifically, barley and maize contain higher levels of phenols, which positively associate with gut microbiome health, oral microbiome health, and overall systemic health [97, 98]. In addition, rice and wheat are digested faster than barley and maize [96]. This higher digestibility might be attributed to differences in cell structure, like thinner cell walls [99]. More starch consumption and more salivary amylase, the first step of starch digestion, may also be contributing factors. Higher salivary amylase copy number has been observed in individuals with high starch diets [100] and is associated with oral microbiome composition [101]. Furthermore, barley has higher fiber content compared to other refined grains [102], which is also associated with improved health outcomes [103]. Finally, barley is reported to have a lower glycemic index compared to rice and wheat [95], which is generally associated with more positive health outcomes [104]. We observe a significant association between grain type and *Brachymonas*, which has not been previously found to be associated with grains in human oral microbiomes. Little is known about the role of this microbe in humans, beyond its presence in healthy oral microbiomes [105–108]. Instead, this microbe has been demonstrated to decrease in abundance in the rumen microbiome of cows fed a high-grain diet [109], so further investigation is needed to fully understand this potential relationship

We also identify a possible relationship between *sisnu* and the oral microbiome. *Sisnu*, also referred to as nettle, is a fibrous plant known for its medicinal benefits and primarily consumed by the Chepang foragers [110]. *Sisnu* consumption is a major differentiating lifestyle factor in this study, as demonstrated by correspondence analysis (**S2 Fig**). Although widely used in traditional medicine, little is known about its role in the microbiome. *Sisnu* shows strong antimicrobial properties against a variety of gram positive and negative bacteria *in vitro* [111], although evidence is mixed [112]. Here, *Porphyromonadaceae_unclassified* relative abundance increases with *sisnu* consumption. Several mechanisms could explain this association. First, ingesting *sisnu* may result in lowered absolute abundance of the oral microbiome overall, with *Porphyromonadaceae_unclassified* being more resilient than other microbes. Second, the antimicrobial effects could create an expanded niche for this microbe to thrive, without impacting absolute abundance levels across the oral microbiome. Finally, the association may be unrelated to potential antimicrobial properties of the plant, but rather nutrition. A finer-grain investigation would be needed to fully establish the underlying causes of this association, especially considering the prominence of *sisnu* as a therapeutic agent. Overall, these analyses of specific lifestyle factors associated with the oral microbiome in Nepali populations provide new insight into the role of specific dietary components and environmental factors in the oral microbiomes of non-industrialized, non-equatorial populations.

One unexpected result was the lack of an observed association with teeth brushing or flossing. Oral hygiene practices are associated with the microbiome, as the mechanical actions of brushing and flossing disrupt plaques and antimicrobial toothpastes also chemically break down biofilms. As a result, even subtle differences in types of toothpaste and brushing frequency result in changes in the plaque and salivary microbiota [113–116]. In traditional populations, miswak, also referred to as a chewing stick, is often utilized for mechanically and chemically cleaning teeth and has been demonstrated to inhibit common oral pathogens [117–121]. Finally, charcoal and ash have also been used historically for oral hygiene, although their effectiveness is highly disputed [122]. Our study included individuals that reportedly do not brush their teeth, those who use traditional methods of miswak or charcoal, and those that brush with a toothbrush and toothpaste. Unexpectedly, we observe no association with reported tooth brushing, let alone brushing method or frequency. This may be due to undocumented variance in “brushers”, as important aspects of oral hygiene such as differing toothpaste amounts, brushing times, and toothpaste type, including whether toothpaste included antibiotics like triclosan, were not fully captured in the administered survey. These factors may have confounded our ability to appropriately identify an association with teeth brushing and the microbiome. In a similar vein, we are also missing insight into oral health, such as disease status or symptoms. Gathering this type of information can only be effectively executed by a dentist, which entails identifying a dental professional willing to go into the field to do so. While challenging, additional data about oral hygiene and oral health would provide insight into how these factors shift with the microbiomes of traditional populations.

Finally, we examined the oral-gut microbiome axis across traditional lifestyles. The oral and gut niches are linked by a consistent one way flow of saliva, food, and microbes from the mouth through to the colon. Although each niche hosts a distinct microbial community of locally adapted strains [123], the microbes originating in the oral cavity can colonize distal sites in the gut and the two communities have been demonstrated to be predictive of one another [124–126]. An estimated one third of oral microbes are able to colonize the guts of healthy individuals [127], with increasing rates of colonization in individuals with diseases such as bowel cancer [128, 129], rheumatoid arthritis [130], and inflammatory bowel diseases [131]. Given the higher incidents of these diseases in industrialized countries [132–134], an open question remains whether rates of translocation along the oral-gut axis vary with lifestyle, potentially being a risk factor for disease. While our 16S rRNA sequence data lacks the resolution needed to detect translocation specifically, we evaluated whether intra-individual similarity between the oral and gut microbiomes increased along the lifestyle gradient from traditional to agrarian. As expected under this hypothesis, there is a decrease in intra-individual Bray-Curtis distances along the gradient of traditional to agrarian populations within Nepal, although not statistically significant. This finding is similar to results from Cameroonian populations, in which the similarity of the oral-gut axis increased across a gradient of rural to urban populations, albeit not significantly [32]. It is worth noting that we did not include Nepali Expats or American Industrialists in this analysis as we did not have paired oral-gut samples for those participants. We also note that the sequencing approach used only provides resolution to the genus level, so we were not able to distinguish whether similarity is a result of translocation of oral strains to the gut versus a homogenization towards similar taxonomic profiles between the niches. Regardless, these results point towards an intriguing hypothesis: oral-gut microbial translocation increases with industrialization, potentially being a risk factor for disease. Future work would entail both increasing sample sizes of individuals across lifestyles as well as using methodologies that reliably track strain sharing across the two body sites.

## Conclusions

Our investigation of Nepali populations across a variety of human lifestyles expands our understanding of the oral microbiome at a global scale. In conjunction with gut microbiomes collected from the same individuals [15], we find that lifestyle is associated with the composition of both the gut and oral microbiomes, albeit to differing degrees. Metagenomic sequencing would provide finer scale microbiome data that would allow us to more effectively identify the taxa and functional potential associated with lifestyle. In addition, studies in which industrialization is decoupled from background genetics, geography, and latitude will be essential for identifying the specific factors that result in microbiome differences across populations. Future work will reveal the extent to which oral microbiomes vary around the globe and refine our understanding of the environmental factors involved.

## Methods

### Ethics approval

This work was approved by the Ethical Review Board of the Nepal Health Research Council (NHRC) as well as by the Stanford University Institutional Review Board. Samples were collected between March and April 2016 with informed consent from all participants.

### Sample Collection

Saliva samples were collected under informed consent using the DNA Genotek Oral Saliva kit (DNA Genotek, Stittsville, CA) from five populations in Nepal: the Chepang (foragers, n = 19), Raji (recently settled/transitioned to agriculture, n = 11), Raute (recently settled/transitioned to agriculture, n = 14), Tharu (established agriculturalist, n = 21), and Newar ethnic groups (established agriculturalist, n = 8). Samples were collected in the winter of 2016 in March and April. Approximately 1mL of unstimulated saliva samples were stored in the stabilizing buffer provided in the DNA Genotek kit and transported over 7-10 days at room temperature, after which they were stored at −20°C for 2-3 months at the Institute of Medicine in Tribhuvan University, Kathmandu, Nepal.

At the time of saliva sample collection, detailed metadata was also collected from all participants. These data include demographic, anthropometric measurements, environmental, and dietary data using a survey questionnaire specifically designed for this study (**S1 Table**). Participants responses to the survey data questionnaires were cleaned and standardized prior to analysis (**S1 Table**).

Following the field work in Nepal, saliva samples were also collected from two populations in the US with the same kits: the Newar population (expats emigrated from Nepal to the US, n = 12) and European Americans (industrialist, n = 6) (Stanford IRB: 35580). USA based samples were collected in the winter of 2016 (November and December). All sampled individuals were unrelated and over 18 years old. Detailed survey data are available for the expat Newar population, but not the European American Industrialists (**S1 Table**). Overall, our cohort included foragers (Chepang, n = 19), recently settled individuals (total, n = 25; Raji, n = 11; Raute, n = 14), established agriculturalists (total, n = 29; Tharu, n = 21; Newar, n = 8), expats (Newar, n = 12), and industrialists (European-Americans, n = 6).

### DNA Extraction and 16S rRNA Amplicon Sequencing

For samples collected in Nepal, total DNA was extracted at the Institute of Medicine (IOM) in Kathmandu using the Qiagen QIAamp MinElute Virus Spin kit (Qiagen, Germantown, MD) according to the manufacturer’s protocol. Both the remaining original saliva samples and the Qiagen kit-extracted DNA were shipped to Stanford University on dry ice and then stored at either −20°C until sequencing (extracted DNA) or at −80℃ (remaining saliva sample). Total DNA was again extracted from saliva samples using the MO BIO PowerSoil DNA Isolation kit (MO BIO, Carlsbad, CA) following the manufacturer’s recommended protocol. For samples collected in the US, they were extracted with both the Qiagen QIAamp MinElute Virus Spin kit and MO BIO PowerSoil DNA Isolation kit in the US. Extraction negative controls were included in all extractions to evaluate contamination during analysis.

The V4 hypervariable region of the 16S rRNA gene was amplified for all DNA extracts and PCR negative controls using established primers and protocols [135]. The sample and negative control libraries were multiplexed and sequenced 250bp single-end on the Illumina MiSeq platform at Stanford University, targeting a minimum of 25,000 reads per sample for accurate relative abundance quantification (**S1 Table**).

### Sequencing Data Quality Control and Cleaning

All bioinformatic analyses were conducted in R version 4.1.2., unless otherwise stated. Single-end sequences were cleaned and processed using DADA2 v.1.22.0 [136]. First, reads were trimmed at 150bp to remove low quality bases, and then filtered to remove any reads with N nucleotides or more than two expected errors (maxN = 0, maxEE = 2, truncQ = 2, **S8 Table**). Next, sequence variants were inferred by pooling reads across samples (pool = TRUE). More specifically, 173 samples were pooled using 8,435,257 reads across 674,656 unique sequences. A sequence table was generated, consisting of 173 samples and 5068 amplicon sequence variants (ASV). 11% of the reads were removed as chimeric, resulting in 7490294 reads and 1424 ASVs remaining (**S8 Table**). ASVs were classified using the RDP v14 training set [137]. Finally, a phylogenetic tree was generated by performing multiple sequence alignment using DECIPHER v.2.22.0 [138] and then constructing the tree by using a neighbor-joining tree as a starting point via the package phangorn v.2.11.1 [139]. ASVs were then handed off to phyloseq v.1.38.0 for additional cleaning and downstream analysis [140].

Next, predicted contaminants were identified and removed via decontam v.1.14.0 by using both the frequency and prevalence methods, thereby removing 19 ASVs [141]. Singletons and any taxa that do not appear at least five times across at least two samples were removed to account for spurious taxa stemming from sequencing errors (**S8 Table**). For alpha diversity analyses, samples were rarefied using the rarefy_even_depth function from the phyloseq package by subsampling to 26923 sequences (the minimum number of sequences across non-control samples) and then calculating alpha diversity from the rarefied samples (**S17 Fig, S18 Fig**). This was repeated 1000 times to account for randomness in rarefaction, and the mean alpha diversity value was calculated. Non-rarefied counts were used for tools with in-built compositionally-aware transformation methods (ie. centered log-ratio transformation). For all other analyses, counts were transformed to relative abundances using total-sum scaling. Log transformations were conducted with log10 and a pseudocount of 0.0001. Finally, individuals currently taking antibiotics were removed (n = 2) (**S19 Fig**), because antibiotics have previously been associated with changes in the oral microbiome [142]. Furthermore, current antibiotic use was found to be marginally significant when compared across Shannon alpha diversity for samples extracted with the Qiagen kit (p = 0.048, Kruskal-Wallis). The post-QC result sample sizes are 60 samples extracted by the PowerSoil kit and 69 samples extracted by the Qiagen kit (**S20 Fig**). The total number of ASVs across all samples extracted via the Qiagen and PowerSoil kits is 1000 ASVs.

### Random Forests Classifier

Random Forests was first conducted by building 500 trees with 63 samples and 37 categorical variables from the survey data using the R package randomForest v.4.7-1.1 [143]. Random Forests models were subsequently conducted with 500 trees using microbiome data agglomerated to the genus level and transformed to relative abundances. Random Forests models were evaluated via out-of-bag error estimates, as well as assessing confusion matrices. Improvement beyond random chance was assessed using the R package verification v.1.42 [144].

### Diversity Analyses

Five metrics were used to assess alpha diversity — Faith’s Phylogenetic Distance, Fisher’s alpha, Shannon alpha, Simpson alpha, and species richness [145–148]. Alpha diversity was calculated after rarefying sample counts to 26923 1000 times and taking the mean value. Kruskal-Wallis tests were performed to assess significant differences in alpha diversity between the ethnic groups. Dunn’s post-hoc test was used to identify the group that was driving the differences.

Beta diversity was calculated from relative abundance counts using Bray-Curtis distance, unweighted Unifrac, and weighted Unifrac [45, 46]. The resulting distances were ordinated using PCoA as implemented in phyloseq. PERMANOVA was performed to assess dissimilarity between ethnic groups using vegan v.2.6-4, permuted 99999 times [149, 150]. Jonckheere-Terpstra tests were used to assess whether the individual PCoA axes followed the lifestyle trend [151].

### Extraction Kit Comparison

The Qiagen QIAamp MinElute Virus Spin and MO BIO PowerSoil DNA Isolation extraction kits were compared based on overall microbiome beta diversity. Beta diversity was calculated and PERMANOVA was performed as described above. Comparisons suggest that while there is some qualitative difference between the two kits, there is no statistically significant difference (PERMANOVA, p > 0.05) (**S3 FigA**), and PCoA axes 1 and 2 are highly correlated across kit (PCoA1 rho = 0.96, p < 2.2*10^-16^; PCoA2 rho = 0.89, p < 2.2*10^-16^) (**S3 FigB**). For fidelity, diversity analyses were conducted using both extraction kits, but other analyses were conducted using only the dataset extracted via the Qiagen kit, selected for its larger sample size (**S3 Fig**).

### Differential Abundance Analysis

Differential abundance analysis was conducted with microbiome count data agglomerated to the genus level using the ALDEx2 v.1.29.2.1. The Kruskal-Wallis module was used to identify microbes differentially abundant across all lifestyles, whereas the standard t-test module was used for assessing microbes differentially abundant across two conditions. To account for multiple tests, the Benjamini-Hochberg method for p-value correction was applied [152]. Effect sizes > 1 and adjusted p-values < 0.05 were considered significant.

To identify which microbes followed the lifestyle trend, Jonckheere-Terpstra tests were performed for each genus using microbiome data agglomerated to the genus level and transformed to relative abundances [151]. All p-values stemming from the Jonckheere-Terpstra tests were corrected for multiple tests using the Benjamini-Hochberg method [152], and adjusted p-values < 0.05 were considered significant.

To validate that the trend observed is not an artifact of sequencing depth, read depth was examined across lifestyle. Sequencing depth was not found to be associated with lifestyle (p > 0.05, Kruskal-Walis; **S21 FigA**). For additional confirmation, *Brachymonas* relative abundance was assessed against read depth, and no correlation was found (p > 0.05, rho = −0.11, Spearman correlation; **S21 FigB**).

### Associations between lifestyle factors, population, and the microbiome

Correspondence Analysis (CA) was performed on the survey metadata using FactoMineR v.2.9 [153]. To determine whether the samples (rows) and lifestyle factors (columns) were significantly associated, a chi-square test of independence was performed. The top 10 most contributing factors for CA1 and CA2 were identified. Alpha and beta diversity values were calculated as described above, and correlations to the CA axes were calculated using Spearman correlation. Canonical Correspondence Analysis (CCA) was performed with microbiome data agglomerated to the genus level, transformed to relative abundances, and then log transformed using vegan v.2.6-4 [150]. The metadata table inputted into CCA was subset to the top 10 categorical lifestyle factors contributing to CA1 and CA2, for a total of 15 factors. Only these 15 factors were used for the CCA model due to multicollinearity between all 37 variables. Model significance and significant lifestyle factors were identified and assessed using the function anova.cca from the vegan package, which performs an ANOVA-like permutation test, permuted 999 times.

To identify which lifestyle factors were specifically associated with a particular taxon, a linear model was generated for the relationship between lifestyle variable and a selected taxon, and then tested for significance. Taxa were agglomerated to the genus level, transformed to relative abundance, and then log transformed for visualization. All p-values stemming from linear models were corrected for multiple tests using the Benjamini-Hochberg method [152], and adjusted p-values < 0.05 were considered significant.

### PICRUSt2 Analysis

PICRUSt2 v2.5.2 was conducted using the “--per_sequence_contrib” option to predict pathway abundances for each ASV [48]. Pathway abundances were predicted using the KEGG database FTP Release 2022-11-07 [154]. Differential abundance analysis and examination of functions following the lifestyle trend were conducted as previously described using the pathway abundance output. To investigate microbiome functional enrichment, differential abundance analysis was conducted with the KEGG Orthologs (KO) and any significant K genes (prior to multiple test correction) were inputted into MicrobiomeProfiler v1.0.0 [155].

### Network Analysis

Network analysis was conducted using the SparCC module in SpiecEasi v.1.1.2 [50, 51]. Microbiome count data agglomerated to the genus level was inputted to generate correlations between taxa. Networks were analyzed for centrality, modularity, and degree distribution using igraph v.1.6.0 [156]. Co-abundance groups (CAGs) were generated using the cluster_fast_greedy function in the igraph package [157]. Networks were plotted in Cytoscape v.3.8.2 [156, 158]. Nodes with 0 edges were removed for visualization and generating CAGs.

### Comparison of the Gut and Oral Microbiomes

The gut and oral microbiomes were only compared between the same individuals overlapping across both studies (**S6 Table**). PCoA axes were generated for the microbiome datasets for each location as previously described and Spearman correlations were calculated between axes. To compare beta diversity between the gut and oral microbiomes within an individual, all microbiome data was combined into one set, Bray Curtis distances were calculated between the gut and oral microbiomes for all samples, and the Bray Curtis distance between the gut and oral microbiomes within an individuals were specifically examined for differences across lifestyle with significance determined by a Jonckheere-Terpstra test.

## Abbreviations

CA: correspondence analysis

PERMANOVA: permutational multivariate analysis of variance using distance matrices

PCoA: Principal Coordinate Analysis

CCA: canonical correspondence analysis

ANOVA: analysis of variance

padj: p-values adjusted for multiple test correction

CAG: co-abundance group

USA: United States of America

IRB: Institutional Review Board

bp: base pair

ASV: amplicon sequence variants

KO: KEGG Orthologs

## Declarations

### Consent for publication

Not applicable.

### Availability of data and materials

Sequence data can be retrieved from the Sequence Read Archive (SRA) under BioProject number PRJNA1098228. All scripts used in this analysis are available at github.com/davenport-lab/Nepali_oral_microbiomes.

### Competing interests

The authors declare that they have no competing interests.

## Funding

PSU-NIH funded Computation, Bioinformatics, and Statistics training grant (T32 GM102057) to EPR. NIH grant R35GM146980 to ERD. Seed Award from Center for Human and Evolutionary Genomics (CEHG, https://cehg.stanford.edu/) and Faculty Research Fund from New York University Abu Dhabi (NYUAD, https://nyuad.nyu.edu/) to ARJ. The Thomas C. and Joan M. Merigan Endowment at Stanford University to DAR. The funders had no role in study design, data collection and analysis, decision to publish, or preparation of the manuscript.

## Authors’ contributions

The study was designed by ARJ and ERD. Samples were collected by YG, ARJ, and GPG. Data generation was performed by DB, ST, and DMP and supervised by JBS and DAR. EPR, AAS, MG, ARJ, and ERD developed the data analysis plan and interpreted results. Computational and statistical analyses were performed by EPR and MG. Finally, EPR, ARJ, and ERD drafted the manuscript. All authors contributed to its revision and approved the final version.

## Acknowledgements

First and foremost, we would like to thank all of the participants of this study. We would also like to thank the Nepal Health Research Council (NHRC) under the Government of the Nepal Ministry of Health for providing research permits to conduct our work in Nepal, as well as Mr. Biswash Chepang and Dil Bhakta Maharjan who acted as community liaisons for the Chepang and Newar communities.

## Additional Files

### Additional file 1 - Supplementary Figures.docx

– Title of data: Supplementary Figures
– Description of data: **S1 Figure**: Confusion matrix of random forest classification based on survey data. **S2 Figure**: Correspondence analysis based on survey data. **S3 Figure**: Microbiome composition within sample is similar across the two extraction kits tested. S4 Figure: All alpha diversity metrics across lifestyle groups with DNA extracted using the Qiagen kit. S5 Figure: Alpha diversity metrics with DNA extracted using the PowerSoil kit. S6 Figure: Oral microbiome composition extended figures - Axis 2 from Qiagen kit extraction, and ordinations on samples extracted using the PowerSoil kit. S7 Figure: Oral microbiome composition calculated with unweighted Unifrac distance. S8 Figure: Oral microbiome composition calculated with weighted Unifrac distance. S9 Figure: Confusion matrix of Random Forest classification based on microbiome data. **S10 Figure**: Correlations between diversity metrics and CA axes. **S11 Figure**: Smoking and *sisnu* are associated with several differentially abundant taxa. **S12 Figure**: Predicted functional abundance significantly differs by lifestyle. **S13 Figure**: Metabolism pathways form the majority of the significantly differentially abundant predicted functions from PICRUSt2. **S14 Figure**: Microbiome functional enrichment analysis. **S15 Figure**: *Fusobacterium* contributes to predicted platinum resistance, which varies by lifestyle. S16 Figure: Correlations between the top three oral and gut microbiome PCoA axes. S17 Figure: Rarefaction curves. S18 Figure: Read depth across samples after read QC. S19 Figure: Comparison of antibiotic use on oral microbiome diversity. S20 Figure: Final sample size per lifestyle per extraction kit. S21 Figure: Read depth against lifestyle and *Brachymonas* relative abundance.

### Additional file 2 - S1 Table.xlsx

– **Title of data:** S1 Table - Sequence, survey, population, and questionnaire info of the sampled individuals
– Description of data: Tab 1 describes survey and sequence metadata data collected. Column abbreviations and responses are explained in greater detail in Tab 3. Column names that end with “2” contain responses that were categorized and transformed to a scale of 0-3, in which possible values for binary variables are 0 or 3 (ie. sex) and possible values for continuous variables are 0, 1, 2, or 3 (ie. fuel source). No survey data was collected for the American Industrialists. Tab 2 describes the lifestyle pertaining to each population and their sample sizes. Tab 3 contains the survey questionnaire, including the codes pertaining to each question asked and list of possible responses.

### Additional file 3 - S2 Table.csv

– Title of data: S2 Table - Oral microbiome differential abundance results from ALDEx2
– Description of data: Oral microbiome genera tested for differential abundance across lifestyle. Overall, 2/111 genera were identified as significantly differentially abundant. Kruskal-Wallis module was utilized and p-value correction was applied using the Benjamini-Hochberg method. Both unadjusted (kw.ep) and adjusted p-values (kw.eBH) are shown, and adjusted p-value < 0.05 is the threshold for significance.

### Additional file 4 - S3 Table.csv

– Title of data: S3 Table - Results of genera tested for following the lifestyle gradient.
– Description of data: All genera tested for following the lifestyle gradient using the Jonckheere-Terpstra test followed by the Benjamini-Hochberg method to correct for multiple tests (BHadj_p_value). Adjusted p-value < 0.05 is the threshold for significance. Nine genera significantly follow the lifestyle gradient.

### Additional file 5 - S4 Table.csv

– Title of data: S4 Table - Associations between differentially abundant microbes and lifestyle factors
– Description of data: Associations between differentially abundant microbes from the Jonckheere-Terpstra test and lifestyle factors were tested via linear models. Linear models were generated between each microbe and each lifestyle factor and then tested for significance, for a total of 333 tested associations. P-value correction was applied using the Benjamini-Hochberg method. Both unadjusted and adjusted p-values are shown. Adjusted p-value < 0.05 is the threshold for significance.

### Additional file 6 - S5 Table.csv

– Title of data: S5 Table - Predicted functional potential differential abundance results
– Description of data: PICRUSt2 predicted functions were analyzed for differential abundance based on lifestyle. None of the 107 tested functions were found to be significant after multiple test correction, but 21/107 pathways were significant prior to correction. Kruskal-Wallis module in ALDEx2 was utilized and p-value correction was applied using the Benjamini-Hochberg method. Both unadjusted (kw.ep) and adjusted p- values (kw.eBH) are shown. Adjusted p-value < 0.05 is the threshold for significance.

### Additional file 7 - S6 Table.csv

– Title of data: S6 Table - Samples overlapping between the gut and oral microbiome studies
– Description of data: List of samples overlapping between the gut and oral microbiome studies, along with the samples unique to each microbiome study. The first column “both” lists sample IDs that are associated with both gut and oral samples. The second column “gut_only” lists sample IDs that are associated with only gut samples. The third column “oral_only” lists sample IDs that are associated with only oral samples.

### Additional file 8 - S7 Table.csv

– Title of data: S7 Table - Gut microbiome differential abundance results from ALDEx2
– Description of data: Gut microbiome genera analyzed for differential abundance based on lifestyle via ALDEx2. Overall, 37/136 genera were identified as significantly differentially abundant. Kruskal-Wallis module was utilized and p-value correction was applied using the Benjamini-Hochberg method. Both unadjusted (kw.ep) and adjusted p- values (kw.eBH) are shown. Adjusted p-value < 0.05 is the threshold for significance.

### Additional file 9 - S8 Table.csv

– Title of data: S8 Table - Read counts through each sequence processing step
– Description of data: Read counts at each step of sequence processing, starting with raw demultiplexed reads and all the way through DADA2, merging, and chimera removal. “Input” column refers to the number of raw reads obtained per sample after sequencing, “filtered” refers to the number of reads remaining after initial read QC, “denoised” refers to the number of reads remaining after denoising in DADA2, “nochim” refers to the number of reads remaining after chimeric sequences were removed, and “retained_overall” is the total proportion of reads retained following all QC steps from the input amount. Table does not include the samples that failed to pass initial read QC.

### Additional file 10 – Nepali abstract.pdf

– Title: Nepali Language Abstract
– Description: Summary of study findings in Nepali language as translated by Aashish R. Jha

## Notes

### Competing Interest Statement

The authors have declared no competing interest.

## References

1. Hublin J-J, Ben-Ncer A, Bailey SE, Freidline SE, Neubauer S, Skinner MM, et al. New fossils from Jebel Irhoud, Morocco and the pan-African origin of Homo sapiens. Nature. 2017;546:289– 92.

2. Alt KW, Al-Ahmad A, Woelber JP. Nutrition and Health in Human Evolution–Past to Present. Nutrients. 2022;14:3594.

3. Clarke SF, Murphy EF, O’Sullivan O, Lucey AJ, Humphreys M, Hogan A, et al. Exercise and associated dietary extremes impact on gut microbial diversity. Gut. 2014;63:1913–20.

4. David LA, Materna AC, Friedman J, Campos-Baptista MI, Blackburn MC, Perrotta A, et al. Host lifestyle affects human microbiota on daily timescales. Genome Biology. 2014;15:R89.

5. Filippo CD, Cavalieri D, Paola MD, Ramazzotti M, Poullet JB, Massart S, et al. Impact of diet in shaping gut microbiota revealed by a comparative study in children from Europe and rural Africa. PNAS. 2010;107:14691–6.

6. Kim S, Covington A, Pamer EG. The intestinal microbiota: Antibiotics, colonization resistance, and enteric pathogens. Immunological Reviews. 2017;279:90–105.

7. Quercia S, Candela M, Giuliani C, Turroni S, Luiselli D, Rampelli S, et al. From lifetime to evolution: timescales of human gut microbiota adaptation. Frontiers in Microbiology. 2014;5.

8. Schnorr SL, Sankaranarayanan K, Lewis CM, Warinner C. Insights into human evolution from ancient and contemporary microbiome studies. Current Opinion in Genetics & Development. 2016;41:14–26.

9. Sonnenburg ED, Sonnenburg JL. The ancestral and industrialized gut microbiota and implications for human health. Nat Rev Microbiol. 2019;17:383–90.

10. Weyrich LS. The evolutionary history of the human oral microbiota and its implications for modern health. Periodontology 2000. 2021;85:90–100.

11. Carter MM, Olm MR, Merrill BD, Dahan D, Tripathi S, Spencer SP, et al. Ultra-deep sequencing of Hadza hunter-gatherers recovers vanishing gut microbes. Cell. 2023;186:3111–3124.e13.

12. Schnorr SL, Candela M, Rampelli S, Centanni M, Consolandi C, Basaglia G, et al. Gut microbiome of the Hadza hunter-gatherers. Nat Commun. 2014;5:3654.

13. Yatsunenko T, Rey FE, Manary MJ, Trehan I, Dominguez-Bello MG, Contreras M, et al. Human gut microbiome viewed across age and geography. Nature. 2012;486:222–7.

14. Gomez A, Petrzelkova KJ, Burns MB, Yeoman CJ, Amato KR, Vlckova K, et al. Gut Microbiome of Coexisting BaAka Pygmies and Bantu Reflects Gradients of Traditional Subsistence Patterns. Cell Reports. 2016;14:2142–53.

15. Jha AR, Davenport ER, Gautam Y, Bhandari D, Tandukar S, Ng KM, et al. Gut microbiome transition across a lifestyle gradient in Himalaya. PLOS Biology. 2018;16:e2005396.

16. Martínez I, Stegen JC, Maldonado-Gómez MX, Eren AM, Siba PM, Greenhill AR, et al. The Gut Microbiota of Rural Papua New Guineans: Composition, Diversity Patterns, and Ecological Processes. Cell Reports. 2015;11:527–38.

17. Baker JL, Mark Welch JL, Kauffman KM, McLean JS, He X. The oral microbiome: diversity, biogeography and human health. Nat Rev Microbiol. 2024;22:89–104.

18. Radaic A, Kapila YL. The oralome and its dysbiosis: New insights into oral microbiome-host interactions. Comput Struct Biotechnol J. 2021;19:1335–60.

19. Willis JR, Gabaldón T. The Human Oral Microbiome in Health and Disease: From Sequences to Ecosystems. Microorganisms. 2020;8:308.

20. Weyrich LS, Dobney K, Cooper A. Ancient DNA analysis of dental calculus. Journal of Human Evolution. 2015;79:119–24.

21. Warinner C. Dental Calculus and the Evolution of the Human Oral Microbiome. J Calif Dent Assoc. 2016;44:411–20.

22. Warinner C, Herbig A, Mann A, Fellows Yates JA, Weiß CL, Burbano HA, et al. A Robust Framework for Microbial Archaeology. Annu Rev Genom Hum Genet. 2017;18:321–56.

23. Adler CJ, Dobney K, Weyrich LS, Kaidonis J, Walker AW, Haak W, et al. Sequencing ancient calcified dental plaque shows changes in oral microbiota with dietary shifts of the Neolithic and Industrial revolutions. Nat Genet. 2013;45:450–5.

24. Gancz AS, Farrer AG, Nixon MP, Wright S, Arriola L, Adler C, et al. Ancient dental calculus reveals oral microbiome shifts associated with lifestyle and disease in Great Britain. Nat Microbiol. 2023;8:2315–25.

25. Ottoni C, Borić D, Cheronet O, Sparacello V, Dori I, Coppa A, et al. Tracking the transition to agriculture in Southern Europe through ancient DNA analysis of dental calculus. PNAS. 2021;118.

26. Quagliariello A, Modi A, Innocenti G, Zaro V, Conati Barbaro C, Ronchitelli A, et al. Ancient oral microbiomes support gradual Neolithic dietary shifts towards agriculture. Nat Commun. 2022;13:6927.

27. Abdill RJ, Adamowicz EM, Blekhman R. Public human microbiome data are dominated by highly developed countries. PLOS Biology. 2022;20:e3001536.

28. Porras AM, Brito IL. The internationalization of human microbiome research. Curr Opin Microbiol. 2019;50:50–5.

29. Araújo V, Fehn A-M, Phiri A, Wills J, Rocha J, Gayà-Vidal M. Oral microbiome homogeneity across diverse human groups from southern Africa: first results from southwestern Angola and Zimbabwe. BMC Microbiol. 2023;23:226.

30. Clemente JC, Pehrsson EC, Blaser MJ, Sandhu K, Gao Z, Wang B, et al. The microbiome of uncontacted Amerindians. Sci Adv. 2015;1:e1500183.

31. Lassalle F, Spagnoletti M, Fumagalli M, Shaw L, Dyble M, Walker C, et al. Oral microbiomes from hunter-gatherers and traditional farmers reveal shifts in commensal balance and pathogen load linked to diet. Molecular Ecology. 2018;27:182–95.

32. Lokmer A, Aflalo S, Amougou N, Lafosse S, Froment A, Tabe FE, et al. Response of the human gut and saliva microbiome to urbanization in Cameroon. Sci Rep. 2020;10:2856.

33. Nasidze I, Li J, Schroeder R, Creasey JL, Li M, Stoneking M. High Diversity of the Saliva Microbiome in Batwa Pygmies. PLOS ONE. 2011;6:e23352.

34. Sprockett DD, Martin M, Costello EK, Burns AR, Holmes SP, Gurven MD, et al. Microbiota assembly, structure, and dynamics among Tsimane horticulturalists of the Bolivian Amazon. Nat Commun. 2020;11:3772.

35. Kidd JM, Sharpton TJ, Bobo D, Norman PJ, Martin AR, Carpenter ML, et al. Exome capture from saliva produces high quality genomic and metagenomic data. BMC Genomics. 2014;15:262.

36. Yeo L-F, Lee SC, Palanisamy UD, Khalid BAK, Ayub Q, Lim SY, et al. The Oral, Gut Microbiota and Cardiometabolic Health of Indigenous Orang Asli Communities. Frontiers in Cellular and Infection Microbiology. 2022;12.

37. Li J, Quinque D, Horz H-P, Li M, Rzhetskaya M, Raff JA, et al. Comparative analysis of the human saliva microbiome from different climate zones: Alaska, Germany, and Africa. BMC Microbiol. 2014;14:316.

38. Ruan X, Luo J, Zhang P, Howell K. The salivary microbiome shows a high prevalence of core bacterial members yet variability across human populations. npj Biofilms Microbiomes. 2022;8:1–14.

39. Thompson LR, Sanders JG, McDonald D, Amir A, Ladau J, Locey KJ, et al. A communal catalogue reveals Earth’s multiscale microbial diversity. Nature. 2017;551:457–63.

40. Fachrul M, Méric G, Inouye M, Pamp SJ, Salim A. Assessing and removing the effect of unwanted technical variations in microbiome data. Sci Rep. 2022;12:22236.

41. Lim MY, Song E-J, Kim SH, Lee J, Nam Y-D. Comparison of DNA extraction methods for human gut microbial community profiling. Systematic and Applied Microbiology. 2018;41:151– 7.

42. Government of Nepal CB of S. National Population and Housing Census 2021. 2021.

43. Kennedy NA, Walker AW, Berry SH, Duncan SH, Farquarson FM, Louis P, et al. The Impact of Different DNA Extraction Kits and Laboratories upon the Assessment of Human Gut Microbiota Composition by 16S rRNA Gene Sequencing. PLOS ONE. 2014;9:e88982.

44. Stinson LF, Keelan JA, Payne MS. Comparison of Meconium DNA Extraction Methods for Use in Microbiome Studies. Front Microbiol. 2018;9.

45. Bray JR, Curtis JT. An Ordination of the Upland Forest Communities of Southern Wisconsin. Ecological Monographs. 1957;27:325–49.

46. Lozupone C, Knight R. UniFrac: a New Phylogenetic Method for Comparing Microbial Communities. Applied and Environmental Microbiology. 2005;71:8228–35.

47. Fernandes AD, Reid JN, Macklaim JM, McMurrough TA, Edgell DR, Gloor GB. Unifying the analysis of high-throughput sequencing datasets: characterizing RNA-seq, 16S rRNA gene sequencing and selective growth experiments by compositional data analysis. Microbiome. 2014;2:15.

48. Douglas GM, Maffei VJ, Zaneveld JR, Yurgel SN, Brown JR, Taylor CM, et al. PICRUSt2 for prediction of metagenome functions. Nat Biotechnol. 2020;38:685–8.

49. Langille MGI. Exploring Linkages between Taxonomic and Functional Profiles of the Human Microbiome. mSystems. 2018;3:e00163–17.

50. Friedman J, Alm EJ. Inferring Correlation Networks from Genomic Survey Data. PLOS Computational Biology. 2012;8:e1002687.

51. Kurtz ZD, Müller CL, Miraldi ER, Littman DR, Blaser MJ, Bonneau RA. Sparse and Compositionally Robust Inference of Microbial Ecological Networks. PLOS Computational Biology. 2015;11:e1004226.

52. Handsley-Davis M, Skelly E, Johnson NW, Kapellas K, Lalloo R, Kroon J, et al. Biocultural Drivers of Salivary Microbiota in Australian Aboriginal and Torres Strait Islander Children. Frontiers in Oral Health. 2021;2.

53. Hansen TH, Kern T, Bak EG, Kashani A, Allin KH, Nielsen T, et al. Impact of a vegan diet on the human salivary microbiota. Sci Rep. 2018;8:5847.

54. Wu J, Peters BA, Dominianni C, Zhang Y, Pei Z, Yang L, et al. Cigarette smoking and the oral microbiome in a large study of American adults. ISME J. 2016;10:2435–46.

55. Yu K-M, Cho H-S, Lee A-M, Lee J-W, Lim S-K. Analysis of the influence of host lifestyle (coffee consumption, drinking, and smoking) on Korean oral microbiome. Forensic Science International: Genetics. 2024;68:102942.

56. Rosas-Plaza S, Hernández-Terán A, Navarro-Díaz M, Escalante AE, Morales-Espinosa R, Cerritos R. Human Gut Microbiome Across Different Lifestyles: From Hunter-Gatherers to Urban Populations. Front Microbiol. 2022;13:843170.

57. Smits SA, Leach J, Sonnenburg ED, Gonzalez CG, Lichtman JS, Reid G, et al. Seasonal Cycling in the Gut Microbiome of the Hadza Hunter-Gatherers of Tanzania. Science. 2017;357:802–6.

58. Blaser MJ. The Past and Future Biology of the Human Microbiome in an Age of Extinctions. Cell. 2018;172:1173–7.

59. Carmody RN, Sarkar A, Reese AT. Gut microbiota through an evolutionary lens. Science. 2021;372:462–3.

60. Ayeni FA, Biagi E, Rampelli S, Fiori J, Soverini M, Audu HJ, et al. Infant and Adult Gut Microbiome and Metabolome in Rural Bassa and Urban Settlers from Nigeria. Cell Reports. 2018;23:3056–67.

61. Obregon-Tito AJ, Tito RY, Metcalf J, Sankaranarayanan K, Clemente JC, Ursell LK, et al. Subsistence strategies in traditional societies distinguish gut microbiomes. Nat Commun. 2015;6:6505.

62. Metcalf CJE, Tepekule B, Bruijning M, Koskella B. Hosts, microbiomes, and the evolution of critical windows. Evolution Letters. 2022;6:412–25.

63. Vangay P, Johnson AJ, Ward TL, Al-Ghalith GA, Shields-Cutler RR, Hillmann BM, et al. US Immigration Westernizes the Human Gut Microbiome. Cell. 2018;175:962–972.e10.

64. Blekhman R, Goodrich JK, Huang K, Sun Q, Bukowski R, Bell JT, et al. Host genetic variation impacts microbiome composition across human body sites. Genome Biology. 2015;16:191.

65. Cameron SJS, Huws SA, Hegarty MJ, Smith DPM, Mur LAJ. The human salivary microbiome exhibits temporal stability in bacterial diversity. FEMS Microbiology Ecology. 2015;91:fiv091.

66. Galloway-Peña JR, Smith DP, Sahasrabhojane P, Wadsworth WD, Fellman BM, Ajami NJ, et al. Characterization of oral and gut microbiome temporal variability in hospitalized cancer patients. Genome Medicine. 2017;9:21.

67. Marotz C, Morton JT, Navarro P, Coker J, Belda-Ferre P, Knight R, et al. Quantifying Live Microbial Load in Human Saliva Samples over Time Reveals Stable Composition and Dynamic Load. mSystems. 2021;6:e01182–20.

68. Zaura E, Brandt BW, Teixeira de Mattos MJ, Buijs MJ, Caspers MPM, Rashid M-U, et al. Same Exposure but Two Radically Different Responses to Antibiotics: Resilience of the Salivary Microbiome versus Long-Term Microbial Shifts in Feces. mBio. 2015;6:e01693–15.

69. Hall MW, Singh N, Ng KF, Lam DK, Goldberg MB, Tenenbaum HC, et al. Inter-personal diversity and temporal dynamics of dental, tongue, and salivary microbiota in the healthy oral cavity. npj Biofilms Microbiomes. 2017;3:1–7.

70. Shi W, Tian J, Xu H, Zhou Q, Qin M. Distinctions and associations between the microbiota of saliva and supragingival plaque of permanent and deciduous teeth. PLoS One. 2018;13:e0200337.

71. Yang X, He L, Yan S, Chen X, Que G. The impact of caries status on supragingival plaque and salivary microbiome in children with mixed dentition: a cross-sectional survey. BMC Oral Health. 2021;21:319.

72. Demirci M. Could Neisseria in oral microbiota modulate the inflammatory response of COVID-19? Oral Dis. 2021;:10.1111/odi.14082.

73. Yamashita Y, Takeshita T. The oral microbiome and human health. J Oral Sci. 2017;59:201– 6.

74. Broussard JL, Devkota S. The changing microbial landscape of Western society: Diet, dwellings and discordance. Mol Metab. 2016;5:737–42.

75. How KY, Song KP, Chan KG. Porphyromonas gingivalis: An Overview of Periodontopathic Pathogen below the Gum Line. Front Microbiol. 2016;7:53.

76. Elliott SP. Rat Bite Fever and Streptobacillus moniliformis. Clin Microbiol Rev. 2007;20:13– 22.

77. Al Bataineh MT, Dash NR, Elkhazendar M, Alnusairat DMH, Darwish IMI, Al-Hajjaj MS, et al. Revealing oral microbiota composition and functionality associated with heavy cigarette smoking. Journal of Translational Medicine. 2020;18:421.

78. Lau SKP, Chan JFW, Tsang C-C, Chan S-M, Ho M-L, Que T-L, et al. Human oropharynx as natural reservoir of Streptobacillus hongkongensis. Sci Rep. 2016;6:24419.

79. Dobon B, Musciotto F, Mira A, Greenacre M, Schlaepfer R, Aguileta G, et al. The making of the oral microbiome in Agta hunter–gatherers. Evolutionary Human Sciences. 2023;5:e13.

80. Aas JA, Griffen AL, Dardis SR, Lee AM, Olsen I, Dewhirst FE, et al. Bacteria of dental caries in primary and permanent teeth in children and young adults. J Clin Microbiol. 2008;46:1407–17.

81. Holgerson PL, Öhman C, Rönnlund A, Johansson I. Maturation of Oral Microbiota in Children with or without Dental Caries. PLOS ONE. 2015;10:e0128534.

82. Obata J, Takeshita T, Shibata Y, Yamanaka W, Unemori M, Akamine A, et al. Identification of the Microbiota in Carious Dentin Lesions Using 16S rRNA Gene Sequencing. PLOS ONE. 2014;9:e103712.

83. Murugesan S, Al Khodor S. Salivary microbiome and hypertension in the Qatari population. Journal of Translational Medicine. 2023;21:454.

84. Alam J, Lee A, Lee J, Kwon DI, Park HK, Park J-H, et al. Dysbiotic oral microbiota and infected salivary glands in Sjögren’s syndrome. PLOS ONE. 2020;15:e0230667.

85. Shouval R, Eshel A, Dubovski B, Kuperman AA, Danylesko I, Fein JA, et al. Patterns of salivary microbiota injury and oral mucositis in recipients of allogeneic hematopoietic stem cell transplantation. Blood Advances. 2020;4:2912–7.

86. Zhou P, Manoil D, Belibasakis GN, Kotsakis GA. Veillonellae: Beyond Bridging Species in Oral Biofilm Ecology. Front Oral Health. 2021;2.

87. Santonocito S, Giudice A, Polizzi A, Troiano G, Merlo EM, Sclafani R, et al. A Cross-Talk between Diet and the Oral Microbiome: Balance of Nutrition on Inflammation and Immune System’s Response during Periodontitis. Nutrients. 2022;14:2426.

88. Maki KA, Ganesan SM, Meeks B, Farmer N, Kazmi N, Barb JJ, et al. The role of the oral microbiome in smoking-related cardiovascular risk: a review of the literature exploring mechanisms and pathways. J Transl Med. 2022;20:584.

89. Shapiro H, Goldenberg K, Ratiner K, Elinav E. Smoking-induced microbial dysbiosis in health and disease. Clinical Science. 2022;136:1371–87.

90. Suzuki N, Nakano Y, Yoneda M, Hirofuji T, Hanioka T. The effects of cigarette smoking on the salivary and tongue microbiome. Clin Exp Dent Res. 2022;8:449–56.

91. Wirth R, Maróti G, Mihók R, Simon-Fiala D, Antal M, Pap B, et al. A case study of salivary microbiome in smokers and non-smokers in Hungary: analysis by shotgun metagenome sequencing. Journal of Oral Microbiology. 2020;12:1773067.

92. Yates JAF, Velsko IM, Aron F, Posth C, Hofman CA, Austin RM, et al. The evolution and changing ecology of the African hominid oral microbiome. PNAS. 2021;118.

93. Klein MI, Duarte S, Xiao J, Mitra S, Foster TH, Koo H. Structural and Molecular Basis of the Role of Starch and Sucrose in Streptococcus mutans Biofilm Development. Applied and Environmental Microbiology. 2009;75:837–41.

94. Saito S, Aoki Y, Tamahara T, Goto M, Matsui H, Kawashima J, et al. Oral Microbiome Analysis in Prospective Genome Cohort Studies of the Tohoku Medical Megabank Project. Frontiers in Cellular and Infection Microbiology. 2021;10.

95. Atkinson FS, Foster-Powell K, Brand-Miller JC. International Tables of Glycemic Index and Glycemic Load Values: 2008. Diabetes Care. 2008;31:2281–3.

96. Soong YY, Tan SP, Leong LP, Henry JK. Total antioxidant capacity and starch digestibility of muffins baked with rice, wheat, oat, corn and barley flour. Food Chemistry. 2014;164:462–9.

97. Esteban-Fernández A, Zorraquín-Peña I, González de Llano D, Bartolomé B, Moreno-Arribas MV. The role of wine and food polyphenols in oral health. Trends in Food Science & Technology. 2017;69:118–30.

98. Narduzzi L, Agulló V, Favari C, Tosi N, Mignogna C, Crozier A, et al. (Poly)phenolic compounds and gut microbiome: new opportunities for personalized nutrition. Microbiome Res Rep. 2022;1:16.

99. Tamura M, Imaizumi R, Saito T, Watanabe T, Okamoto T. Studies of the texture, functional components and *in vitro* starch digestibility of rolled barley. Food Chemistry. 2019;274:672–8.

100. Perry GH, Dominy NJ, Claw KG, Lee AS, Fiegler H, Redon R, et al. Diet and the evolution of human amylase gene copy number variation. Nat Genet. 2007;39:1256–60.

101. Poole AC, Goodrich JK, Youngblut ND, Luque GG, Ruaud A, Sutter JL, et al. Human Salivary Amylase Gene Copy Number Impacts Oral and Gut Microbiomes. Cell Host & Microbe. 2019;25:553–564.e7.

102. Wursch P, Pi-Sunyer FX. The Role of Viscous Soluble Fiber in the Metabolic Control of Diabetes. DIABETES CARE. 1997;20.

103. Li J, Kaneko T, Qin L-Q, Wang J, Wang Y, Sato A. Long-term effects of high dietary fiber intake on glucose tolerance and lipid metabolism in GK rats: comparison among barley, rice, and cornstarch. Metabolism. 2003;52:1206–10.

104. Jenkins DJ, Kendall CW, Augustin LS, Franceschi S, Hamidi M, Marchie A, et al. Glycemic index: overview of implications in health and disease. The American Journal of Clinical Nutrition. 2002;76:266S–273S.

105. Liu S, Xie G, Chen M, He Y, Yu W, Chen X, et al. Oral microbial dysbiosis in patients with periodontitis and chronic obstructive pulmonary disease. Front Cell Infect Microbiol. 2023;13.

106. Liu Y, Liu H, Rong Y, Shi Q, Yang Q, Li H, et al. Alterations of oral microbiota are associated with the development and severity of acute pancreatitis. Journal of Oral Microbiology. 2023;15:2264619.

107. Manzoor M, Lommi S, Furuholm J, Sarkkola C, Engberg E, Raju S, et al. High abundance of sugar metabolisers in saliva of children with caries. Sci Rep. 2021;11:4424.

108. Raju SC, Lagström S, Ellonen P, de Vos WM, Eriksson JG, Weiderpass E, et al. Gender-Specific Associations Between Saliva Microbiota and Body Size. Front Microbiol. 2019;10:767.

109. Petri RM, Wetzels SU, Qumar M, Khiaosa-ard R, Zebeli Q. Adaptive responses in short-chain fatty acid absorption, gene expression, and bacterial community of the bovine rumen epithelium recovered from a continuous or transient high-grain feeding. Journal of Dairy Science. 2019;102:5361–78.

110. Sapkota S, Shrestha S. An explorative survey on Sisnu: A wonder but highly underutilized crop of Nepal. J Pharmacogn Phytochem. 2018;7:832–3.

111. Kregiel D, Pawlikowska E, Antolak H. Urtica spp.: Ordinary Plants with Extraordinary Properties. Molecules. 2018;23:1664.

112. Harrison F, Furner-Pardoe J, Connelly E. An assessment of the evidence for antibacterial activity of stinging nettle (Urtica dioica) extracts. Access Microbiol. 2022;4:000336.

113. Carda-Diéguez M, Moazzez R, Mira A. Functional changes in the oral microbiome after use of fluoride and arginine containing dentifrices: a metagenomic and metatranscriptomic study. Microbiome. 2022;10:159.

114. Adams SE, Arnold D, Murphy B, Carroll P, Green AK, Smith AM, et al. A randomised clinical study to determine the effect of a toothpaste containing enzymes and proteins on plaque oral microbiome ecology. Sci Rep. 2017;7:43344.

115. Kong J, Zhang G, Xia K, Diao C, Yang X, Zuo X, et al. Tooth brushing using toothpaste containing theaflavins reduces the oral pathogenic bacteria in healthy adults. 3 Biotech. 2021;11:150.

116. Calderon SJ, Chung SY, Fields CJ, Mortimer NT. Children Tooth Brushing Behavior and Oral Microbiota: A Pilot Study. Oral. 2021;1:112–21.

117. Sukkarwalla A, Ali SM, Lundberg P, Tanwir F. Efficacy of Miswak on Oral Pathogens. Dent Res J (Isfahan). 2013;10:314–20.

118. Sofrata AH, Claesson RLK, Lingström PK, Gustafsson AK. Strong Antibacterial Effect of Miswak Against Oral Microorganisms Associated With Periodontitis and Caries. Journal of Periodontology. 2008;79:1474–9.

119. Abhary M, Al-Hazmi A-A. Antibacterial activity of Miswak (*Salvadora persica* L.) extracts on oral hygiene. Journal of Taibah University for Science. 2016;10:513–20.

120. Rifaey N, AlAdwani M, Karched M, Baskaradoss JK. A clinical investigation into the efficacy of miswak chewing sticks as an oral hygiene aid: A crossover randomized trial. International Journal of Dental Hygiene. 2021;19:223–30.

121. Sofrata A, Brito F, Al-Otaibi M, Gustafsson A. Short term clinical effect of active and inactive *Salvadora persica* miswak on dental plaque and gingivitis. Journal of Ethnopharmacology. 2011;137:1130–4.

122. Brooks JK, Bashirelahi N, Reynolds MA. Charcoal and charcoal-based dentifrices: A literature review. The Journal of the American Dental Association. 2017;148:661–70.

123. Lloyd-Price J, Mahurkar A, Rahnavard G, Crabtree J, Orvis J, Hall AB, et al. Strains, functions and dynamics in the expanded Human Microbiome Project. Nature. 2017;550:61–6.

124. Ding T, Schloss PD. Dynamics and associations of microbial community types across the human body. Nature. 2014;509:357–60.

125. Kageyama S, Sakata S, Ma J, Asakawa M, Takeshita T, Furuta M, et al. High-Resolution Detection of Translocation of Oral Bacteria to the Gut. J Dent Res. 2023;102:752–8.

126. Liao C, Rolling T, Djukovic A, Fei T, Mishra V, Liu H, et al. Oral bacteria relative abundance in faeces increases due to gut microbiota depletion and is linked with patient outcomes. Nat Microbiol. 2024;9:1555–65.

127. Schmidt TS, Hayward MR, Coelho LP, Li SS, Costea PI, Voigt AY, et al. Extensive transmission of microbes along the gastrointestinal tract. eLife. 2019;8:e42693.

128. Flynn KJ, Baxter NT, Schloss PD. Metabolic and Community Synergy of Oral Bacteria in Colorectal Cancer. mSphere. 2016;1:10.1128/msphere.00102-16.

129. Zeller G, Tap J, Voigt AY, Sunagawa S, Kultima JR, Costea PI, et al. Potential of fecal microbiota for early-stage detection of colorectal cancer. Molecular Systems Biology. 2014;10:766.

130. Zhang X, Zhang D, Jia H, Feng Q, Wang D, Liang D, et al. The oral and gut microbiomes are perturbed in rheumatoid arthritis and partly normalized after treatment. Nat Med. 2015;21:895–905.

131. Gevers D, Kugathasan S, Denson LA, Vázquez-Baeza Y, Van Treuren W, Ren B, et al. The Treatment-Naive Microbiome in New-Onset Crohn’s Disease. Cell Host & Microbe. 2014;15:382–92.

132. Arnaout AY, Nerabani Y, Douba Z, Kassem LH, Arnaout K, Shabouk MB, et al. The prevalence and risk factors of irritable bowel syndrome (PRIBS study) among adults in low- and middle-income countries: A multicenter cross-sectional study. Health Science Reports. 2023;6:e1592.

133. Bray F, Ferlay J, Soerjomataram I, Siegel RL, Torre LA, Jemal A. Global cancer statistics 2018: GLOBOCAN estimates of incidence and mortality worldwide for 36 cancers in 185 countries. CA Cancer J Clin. 2018;68:394–424.

134. GBD 2021 Rheumatoid Arthritis Collaborators. Global, regional, and national burden of rheumatoid arthritis, 1990–2020, and projections to 2050: a systematic analysis of the Global Burden of Disease Study 2021. Lancet Rheumatol. 2023;5:e594–610.

135. Caporaso JG, Lauber CL, Walters WA, Berg-Lyons D, Huntley J, Fierer N, et al. Ultra-high-throughput microbial community analysis on the Illumina HiSeq and MiSeq platforms. ISME J. 2012;6:1621–4.

136. Callahan BJ, McMurdie PJ, Rosen MJ, Han AW, Johnson AJA, Holmes SP. DADA2: High-resolution sample inference from Illumina amplicon data. Nature Methods. 2016;13:581–3.

137. Cole JR, Wang Q, Cardenas E, Fish J, Chai B, Farris RJ, et al. The Ribosomal Database Project: improved alignments and new tools for rRNA analysis. Nucleic Acids Research. 2009;37 suppl_1:D141–5.

138. Wright E S. Using DECIPHER v2.0 to Analyze Big Biological Sequence Data in R. The R Journal. 2016;8:352.

139. Schliep KP. phangorn: phylogenetic analysis in R. Bioinformatics. 2011;27:592–3.

140. McMurdie PJ, Holmes S. phyloseq: An R Package for Reproducible Interactive Analysis and Graphics of Microbiome Census Data. PLOS ONE. 2013;8:e61217.

141. Davis NM, Proctor DM, Holmes SP, Relman DA, Callahan BJ. Simple statistical identification and removal of contaminant sequences in marker-gene and metagenomics data. Microbiome. 2018;6:226.

142. Cheng X, He F, Si M, Sun P, Chen Q. Effects of Antibiotic Use on Saliva Antibody Content and Oral Microbiota in Sprague Dawley Rats. Frontiers in Cellular and Infection Microbiology. 2022;12.

143. Liaw A, Wiener M. Classification and Regression by randomForest. 2002;2.

144. Laboratory N-RA. verification: Weather Forecast Verification Utilities. 2015.

145. Faith DP. Conservation evaluation and phylogenetic diversity. Biological Conservation. 1992;61:1–10.

146. Fisher RA, Corbet AS, Williams CB. The Relation Between the Number of Species and the Number of Individuals in a Random Sample of an Animal Population. Journal of Animal Ecology. 1943;12:42–58.

147. Shannon CE. A Mathematical Theory of Communication. Bell System Technical Journal. 1948;27:379–423.

148. Simpson EH. Measurement of Diversity. Nature. 1949;163:688–688.

149. Anderson MJ. Permutational Multivariate Analysis of Variance (PERMANOVA). In: Wiley StatsRef: Statistics Reference Online. John Wiley & Sons, Ltd; 2017. p. 1–15.

150. Dixon P. VEGAN, a package of R functions for community ecology. Journal of Vegetation Science. 2003;14:927–30.

151. Lunneborg CE. Jonckheere–Terpstra Test. In: Wiley StatsRef: Statistics Reference Online. John Wiley & Sons, Ltd; 2014.

152. Benjamini Y, Hochberg Y. Controlling the False Discovery Rate: A Practical and Powerful Approach to Multiple Testing. Journal of the Royal Statistical Society Series B (Methodological). 1995;57:289–300.

153. Lê S, Josse J, Husson F. FactoMineR: An R Package for Multivariate Analysis. Journal of Statistical Software. 2008;25:1–18.

154. Kanehisa M, Goto S. KEGG: kyoto encyclopedia of genes and genomes. Nucleic Acids Res. 2000;28:27–30.

155. Chen M, Yu G. MicrobiomeProfiler: An R/shiny package for microbiome functional enrichment analysis. 2023.

156. Csardi G, Nepusz T. The Igraph Software Package for Complex Network Research. InterJournal. 2005;Complex Systems:1695.

157. Clauset A, Newman MEJ, Moore C. Finding community structure in very large networks. Phys Rev E. 2004;70:066111.

158. Shannon P, Markiel A, Ozier O, Baliga NS, Wang JT, Ramage D, et al. Cytoscape: A Software Environment for Integrated Models of Biomolecular Interaction Networks. Genome Res. 2003;13:2498–504.

