## Supplemental figures for "Nepali oral microbiomes reflect a gradient of lifestyles from traditional to industrialized"

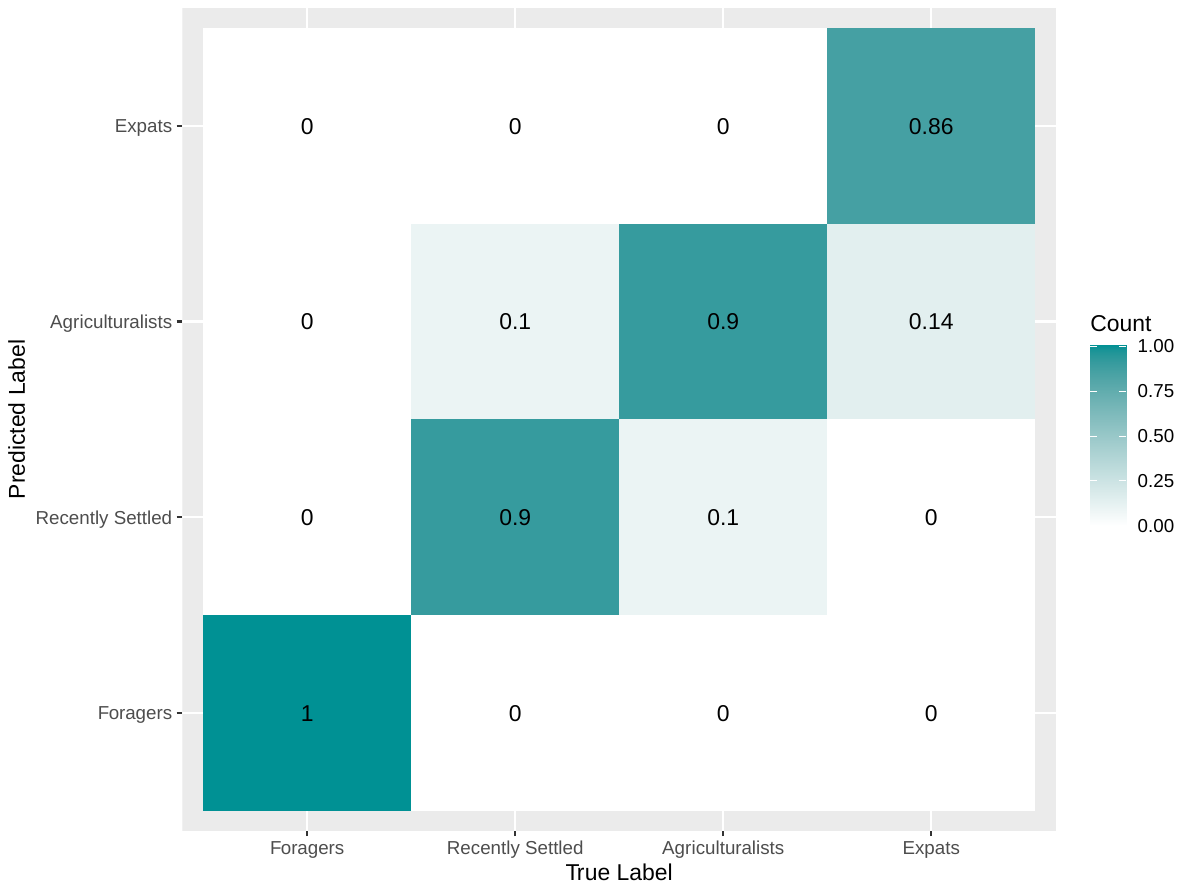

**S1 Figure: Confusion matrix of random forest classification based on survey data**

Confusion matrix showing the results of Random Forests categorizing individuals into their respective lifestyles based on survey data. Random Forests were conducted by building 500 trees with survey data pertaining to all Nepali individuals. American Industrialists were not included due to lack of survey data, so individuals were classified as Forager, Recently Settled, Agriculturalist, or Expat. Any variables that were not collected across all Nepali individuals were removed, resulting in 63 samples and 37 categorical variables from the survey data being used for Random Forest classification. The confusion matrix shows whether individuals were correctly labeled, and if incorrectly labeled, what they were labeled as. The x-axis shows the true lifestyle label of the individual, whereas the y-axis shows the predicted label. The scale ranges from 0 to 1, in which 1 means 100% of the individuals with a particular true label were categorized as a particular predicted lifestyle, whereas 0 means 0% of the individuals with a true label were categorized as a specific lifestyle. The confusion matrix shows that most individuals were correctly categorized as their true lifestyle, with a few individuals being mistakenly categorized as Agriculturalists or Recently settled.

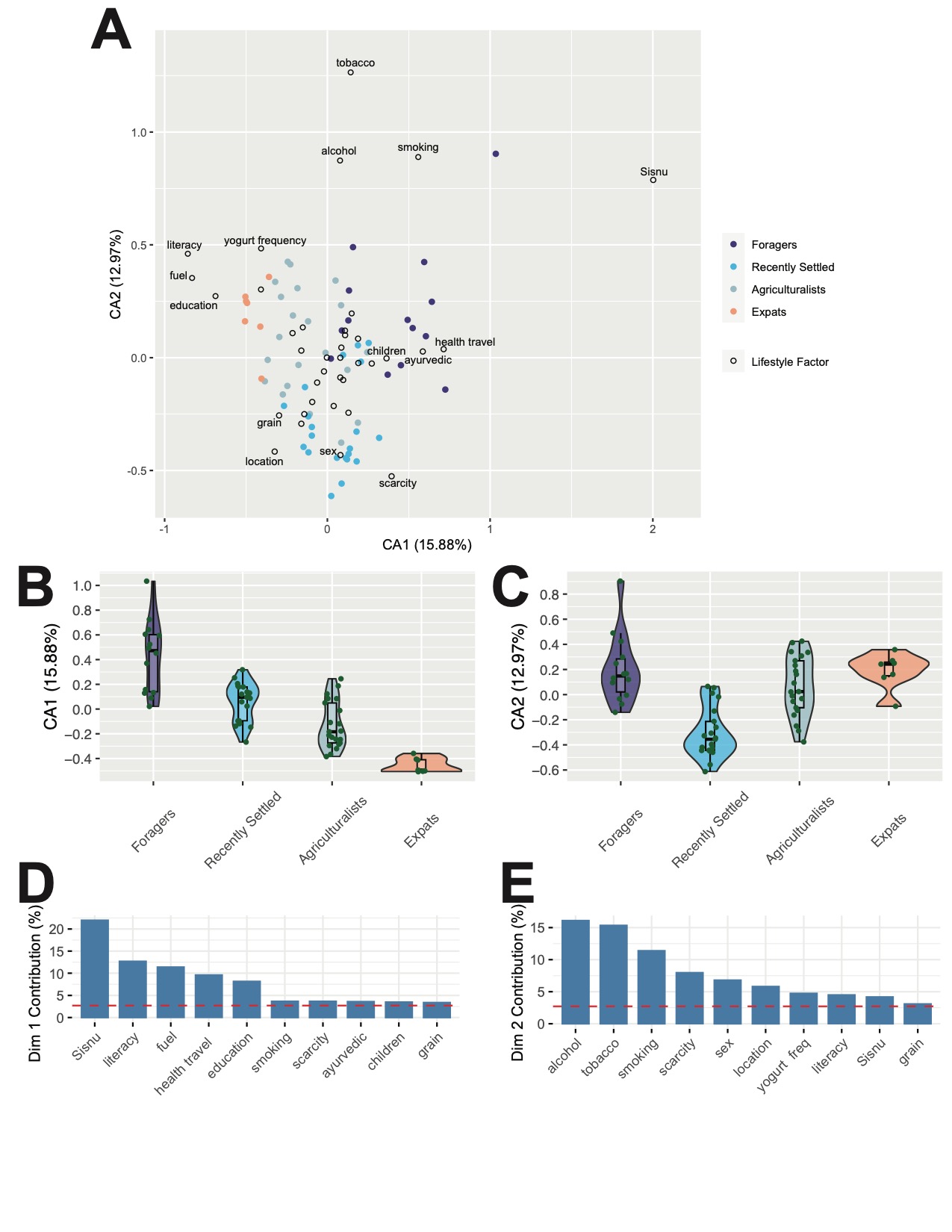

**S2 Figure: Correspondence analysis based on survey data**

A) Symmetric CA biplot representing the lifestyle factors and individuals in the same space. Individuals are colored by lifestyle and factors are unfilled points. Only distance within factors or within individuals can be interpreted; not distance between factor and individual. The top 15 contributing lifestyle factors are labeled. In general, individuals cluster closely, and similar lifestyle factors also cluster closely to each other. B) Distribution of individuals along CA axis 1, grouped by lifestyle. The pattern of individuals follows the lifestyle gradient when examined based on lifestyle factor. C) Distribution of individuals along CA axis 2, grouped by lifestyle. No trend following the lifestyle gradient is observed. D) Top 10 lifestyle factors that contribute most to CA axis 1. The dotted line refers to expected contribution value if all variables contributed equally. E) Top 10 lifestyle factors that contribute most to CA axis 2. Several factors overlap between CA axes 1 and 2, so a total of 15 lifestyle factors are highlighted in part A. The dotted line refers to expected contribution value if all variables contributed equally.

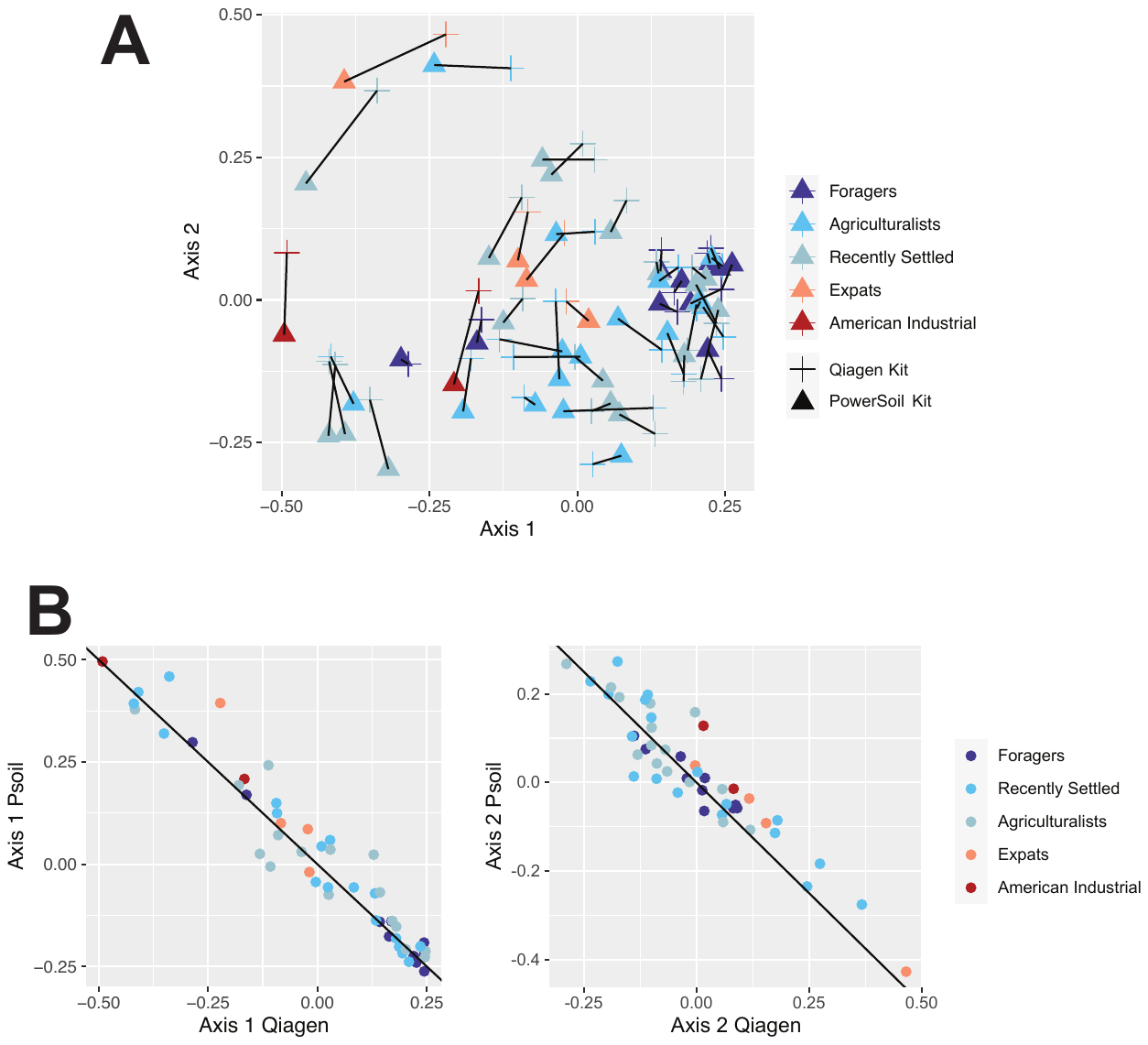

**S3 Figure: Microbiome composition within sample is similar across the two extraction kits tested**

The Qiagen QIAamp MinElute Virus Spin and MO BIO PowerSoil DNA Isolation kits were both used to extract the microbiome from saliva samples. A) PCoA plot with both kits depicted by point shape (Qiagen = plus, PowerSoil = triangle; Bray Curtis dissimilarity). Points are colored by lifestyle and points from the same individual across kits are connected by a line. There is no statistically significant difference between the two kits (PERMANOVA, p > 0.05). B) The first and second PCoA axes values are highly correlated for samples extracted by each kit (PCoA1 rho = 0.96, p < 2.2*10^-16^; PCoA2 rho = 0.89, p < 2.2*10^-16^). Points are colored by lifestyle and black line indicates perfect overlap between the kits.

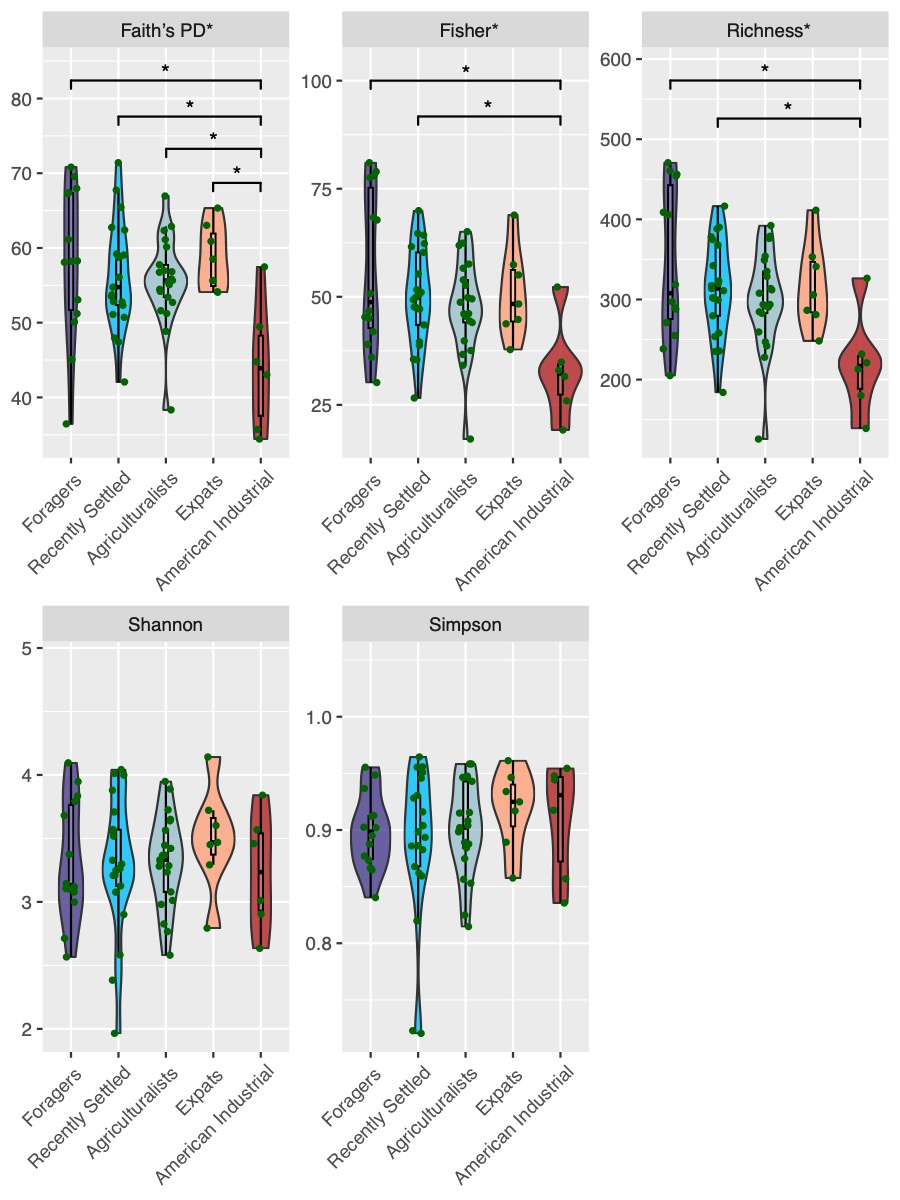

**S4 Figure: All alpha diversity metrics across lifestyle groups with DNA extracted using the Qiagen kit.**

All alpha diversity metrics tested are shown for all individuals, grouped by lifestyle. Data presented here was all extracted using the Qiagen kit. Marginal to no significant differences are observed across all metrics, showing that alpha diversity is consistently not significant across all Nepali individuals regardless of metric. Shannon alpha (bottom left) and Simpson’s diversity index (bottom middle) are not significant (p = 0.8, 0.73; respectively, Kruskal-Wallis). Faith’s phylogenetic diversity (Faith’s PD - top left), Fisher’s alpha diversity (top middle), species richness (top right) are marginally significant (p = 0.028, 0.046, 0.046; respectively, Kruskal-Wallis), as indicated by *.

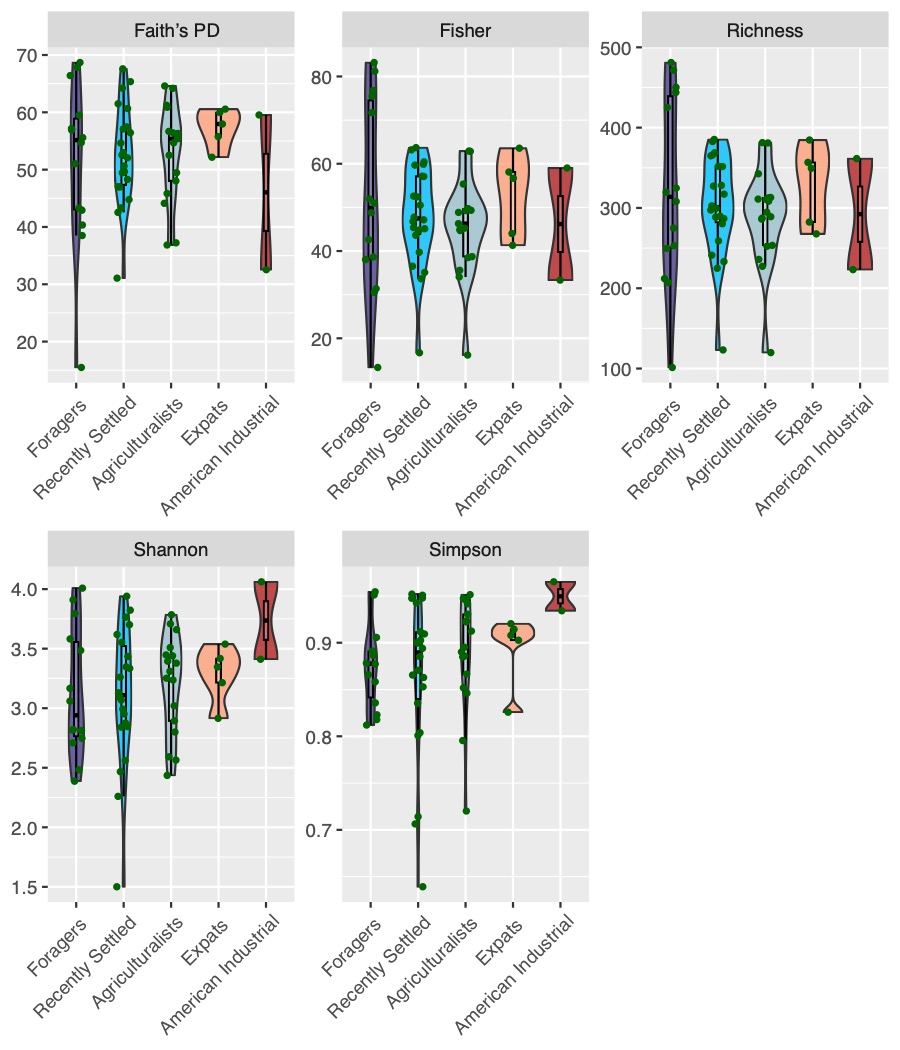

**S5 Figure: Alpha diversity metrics with DNA extracted using the PowerSoil kit.**

All alpha diversity metrics calculated across all individuals using data extracted via the PowerSoil kit. All metrics do not significantly differ based on lifestyle (p > 0.05, Kruskal-Wallis). Alpha diversity findings remain consistent across both extraction kits tested. See Methods for additional details regarding the extraction kit comparison.

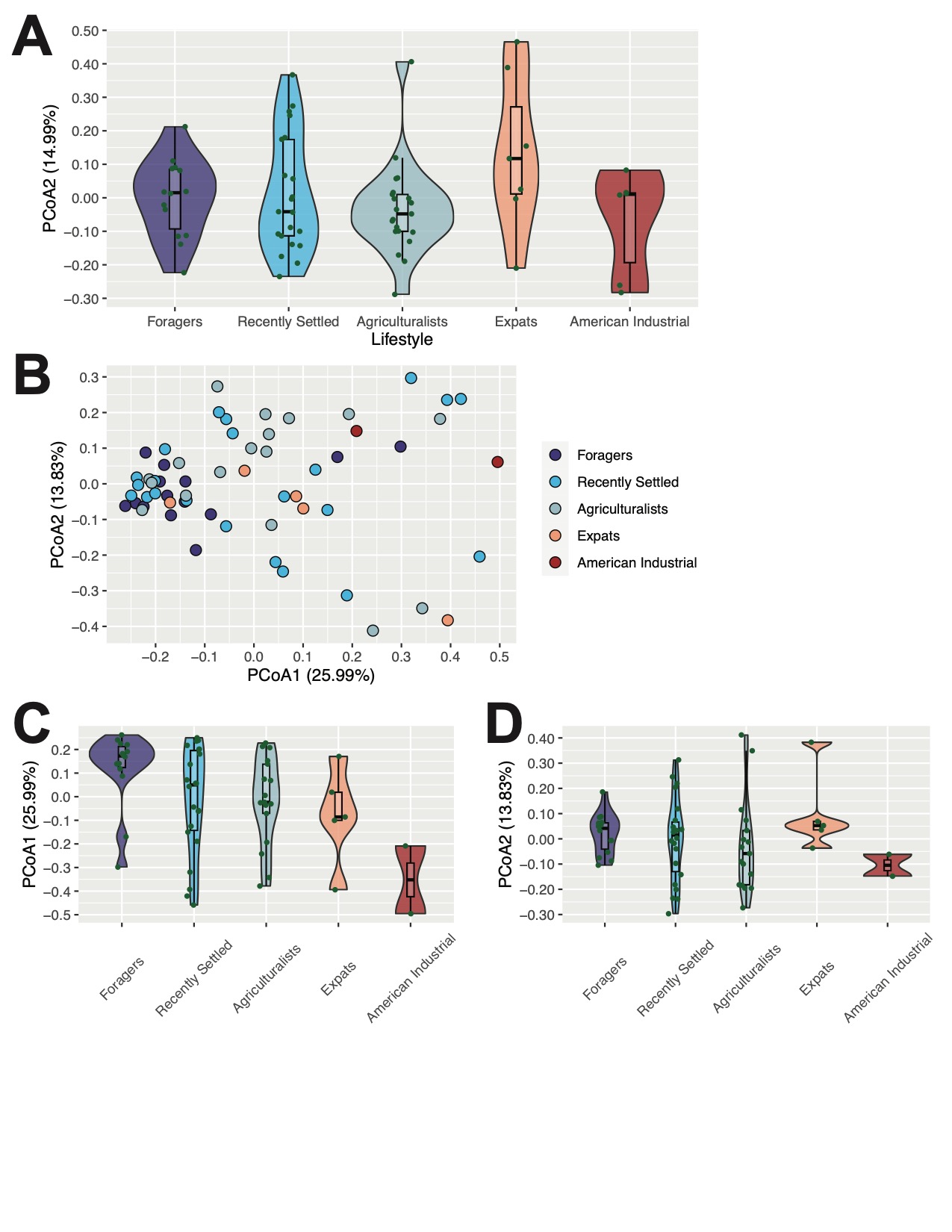

**S6 Figure: Oral microbiome composition extended figures - Axis 2 from Qiagen kit extraction, and ordinations on samples extracted using the PowerSoil kit.**

A) Distribution of individuals along PCoA axis 2, grouped by lifestyle. DNA was extracted with the Qiagen kit as in Figure 3. The distribution does not follow the lifestyle gradient (p > 0.05, Jonckheere-Terpstra test). B) PCoA plot showing individuals ordinated based on Bray-Curtis distance and colored by lifestyle. DNA was extracted via the PowerSoil kit. Overall microbiome diversity varies significantly with lifestyle (p= 0.0081, PERMANOVA). C) Distribution of individuals along PCoA axis 1, grouped by lifestyle. Data was extracted via the PowerSoil kit. The distribution of individuals along PCoA axis 1 follows the lifestyle gradient (p = 0.0039, Jonckheere-Terpstra test). D) Distribution of individuals along PCoA axis 2, grouped by lifestyle. Data was extracted via the PowerSoil kit. The distribution of individuals along PCoA axis 2 does not follow the lifestyle gradient (p = 0.36, Jonckheere-Terpstra test). Beta diversity findings remain consistent across both extraction kits tested. See Methods for additional details regarding the extraction kit comparison.

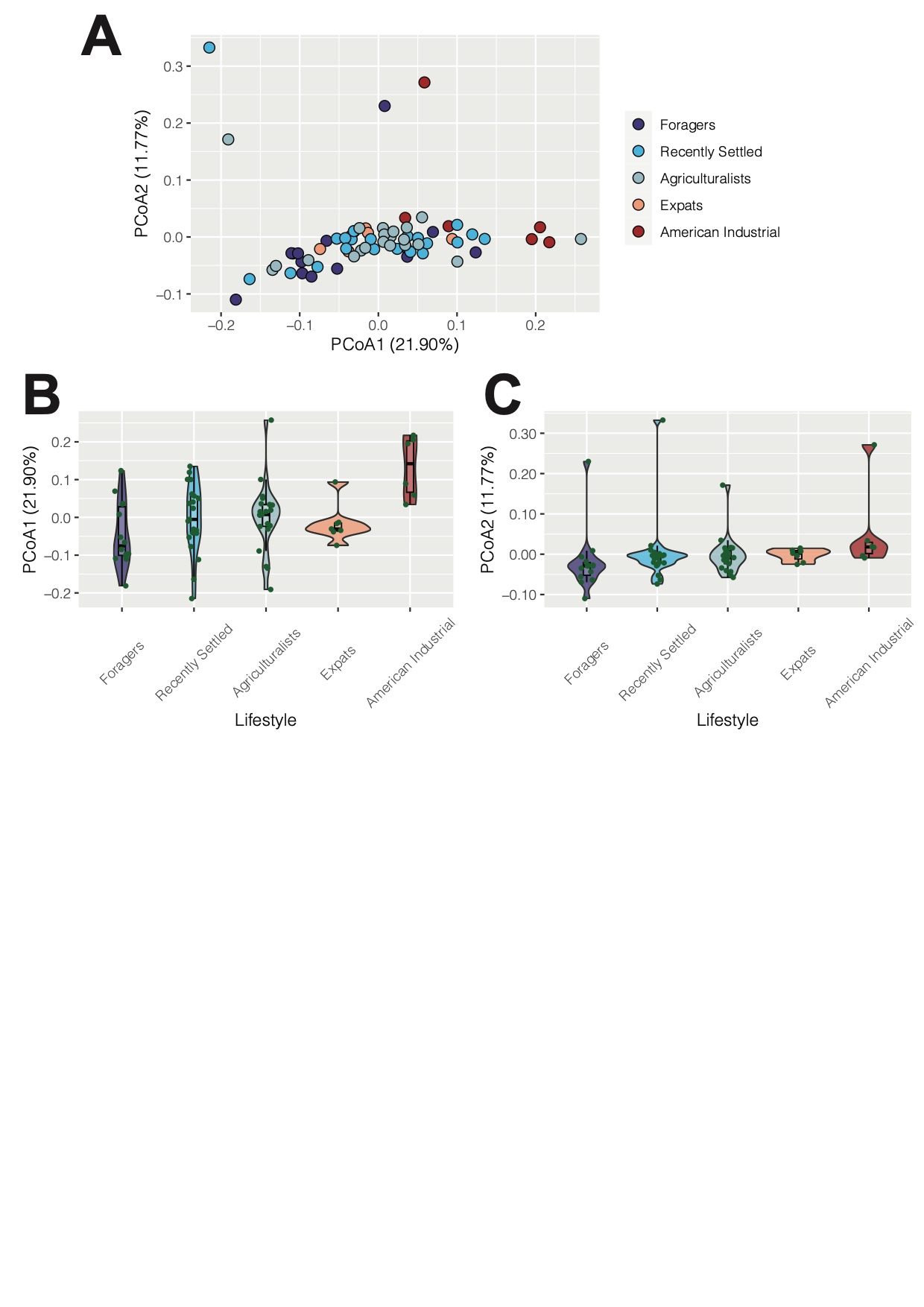

**S7 Figure: Oral microbiome composition calculated with unweighted Unifrac distance**

A) Microbiome composition varies significantly with lifestyle (p = 1x10^-5^, PERMANOVA). The PCoA plot shows individuals ordinated based on unweighted Unifrac distance and colored by lifestyle. B) The distribution of individuals along PCoA axis 1 follows the lifestyle gradient, from traditional to Industrial (p = 0.024, Jonckheere-Terpstra test). Lifestyles are ordered from most traditional (Foragers) to most industrialized (American Industrialists), left to right. C) Distribution of individuals along PCoA axis 2, grouped by lifestyle. The distribution follows the lifestyle gradient (p = 3.5x10^-4^, Jonckheere-Terpstra test).

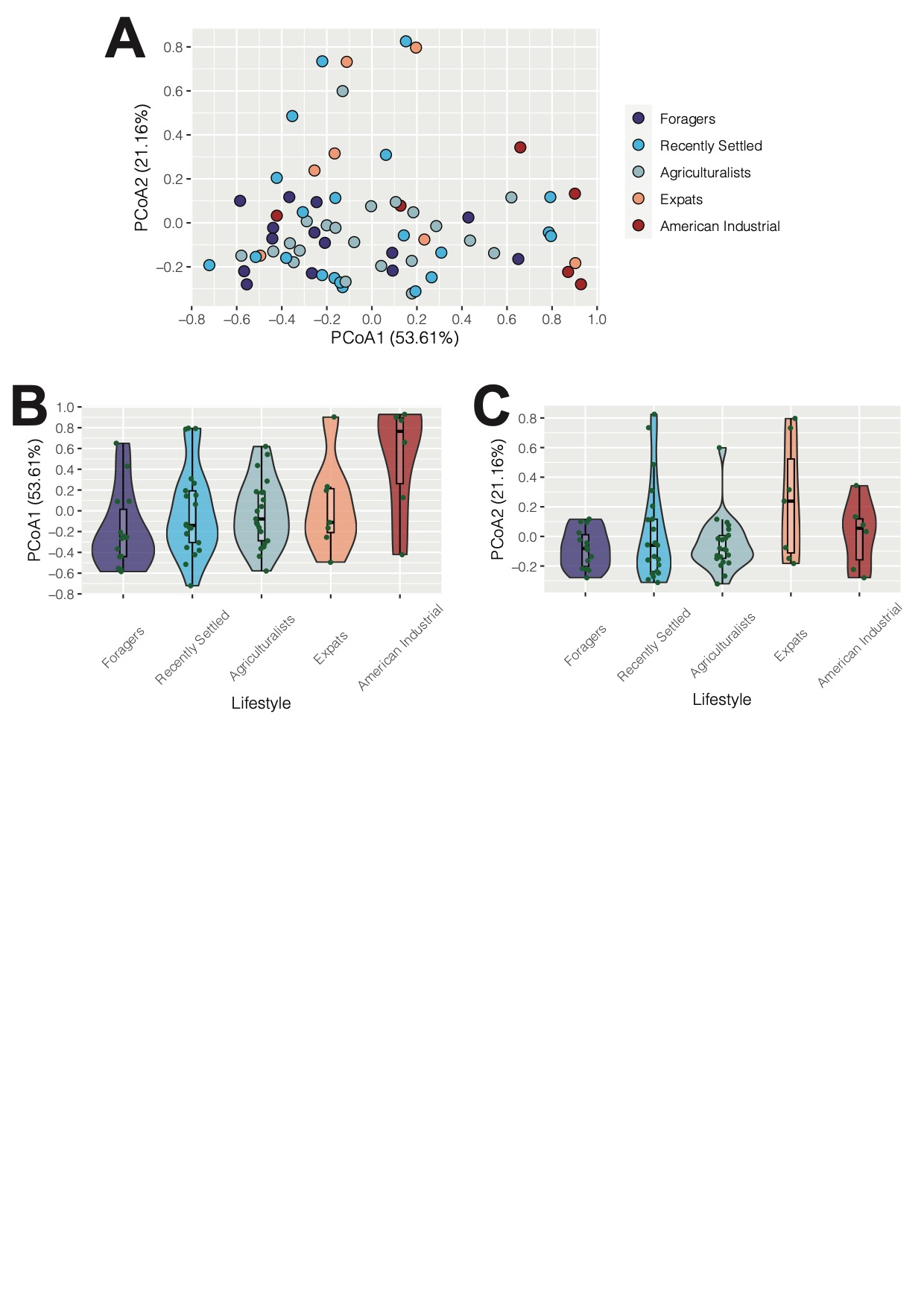

**S8 Figure: Oral microbiome composition calculated with weighted Unifrac distance**

A) Microbiome composition varies significantly with lifestyle (p = 0.0096, PERMANOVA). The PCoA plot shows individuals ordinated based on unweighted Unifrac distance and colored by lifestyle. B) The distribution of individuals along PCoA axis 1 follows the lifestyle gradient, from traditional to Industrial (p = 0.006, Jonckheere-Terpstra test). Lifestyles are ordered from most traditional (Foragers) to most industrialized (American Industrialists), left to right. C) Distribution of individuals along PCoA axis 2, grouped by lifestyle. The distribution does not follow the lifestyle gradient (p > 0.05, Jonckheere-Terpstra test).

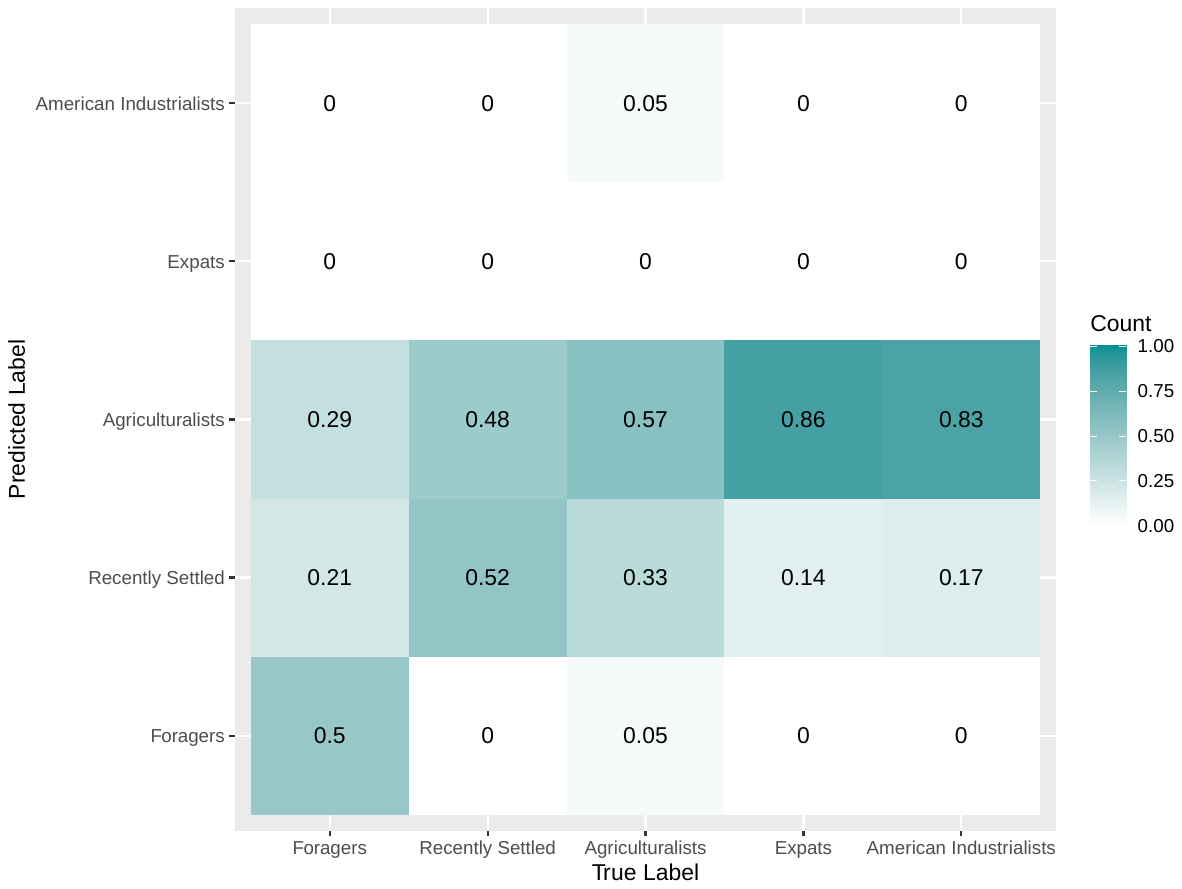

**S9 Figure: Confusion matrix of Random Forest classification based on microbiome data**

Confusion matrix showing the results of Random Forests categorizing individuals into their respective lifestyle groupings based on microbiome data. Random Forests were conducted by building 500 trees with the microbiome data, and individuals were classified as Forager, Recently Settled, Agriculturalist, Expats, or American Industrialists. A total of 69 samples and 1000 ASVs were used for Random Forest classification. The confusion matrix shows whether individuals were correctly labeled, and if incorrectly labeled, what they were labeled as. The x-axis shows the true lifestyle label of the individual, whereas the y-axis shows the predicted label. The scale ranges from 0 to 1, in which 1 means 100% of the individuals with a particular true label were categorized as a particular predicted lifestyle, whereas 0 means 0% of the individuals with a true label were categorized as a specific lifestyle. The confusion matrix shows that most individuals were classified as either Agriculturalist or Recently Settled, with an overall accuracy of 43.48% with a Heidke skill score (HSS) of 0.062. HSS assesses the accuracy of a random forest compared to random chance. Scores range from -1 to 1, where 1 indicates perfect accuracy whereas 0 indicates no skill. HSS > 0.3 indicates positive improvement over random chance. Because the HSS for this random forest is below the threshold, the random forest does not perform better than random chance.

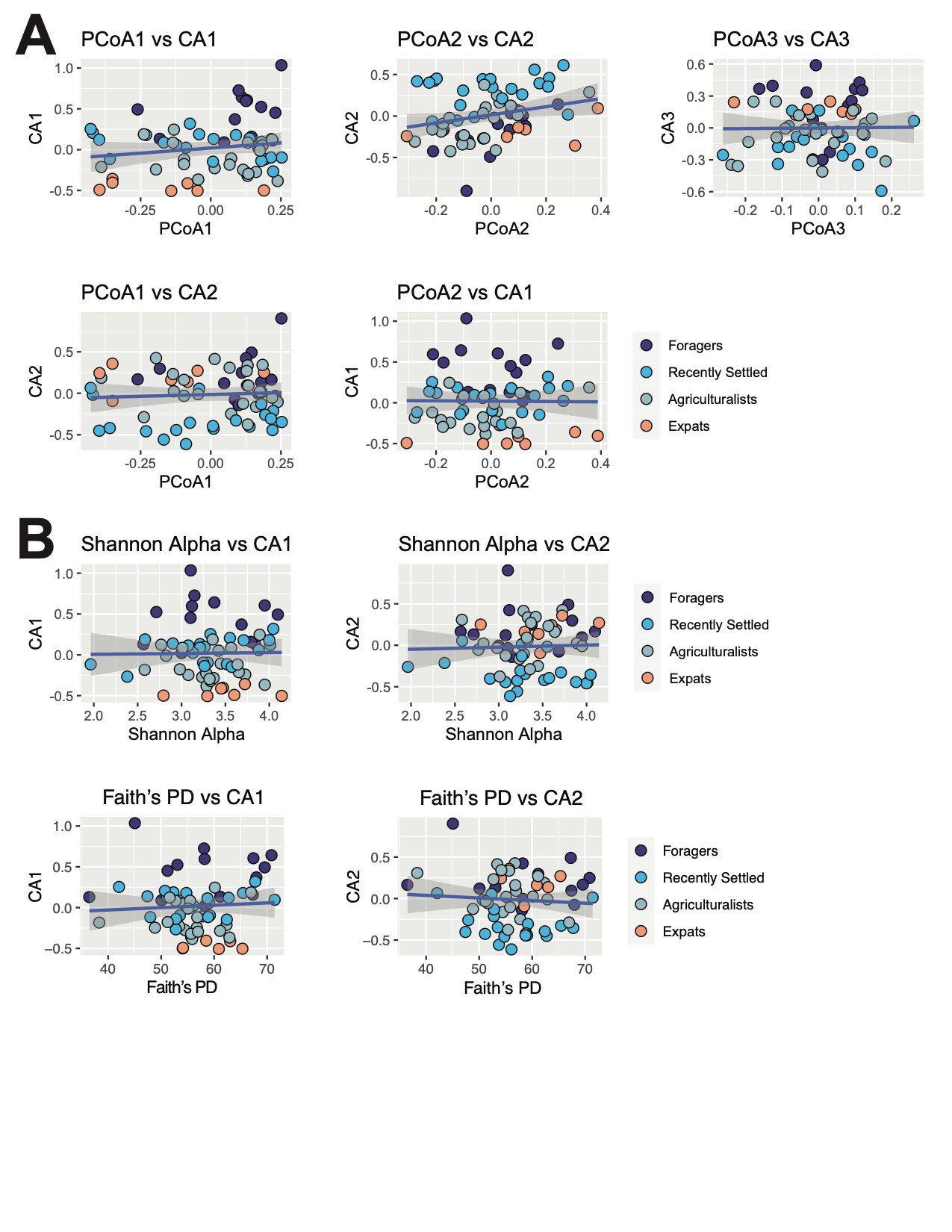

**S10 Figure: Correlations between diversity metrics and CA axes**

A) Correlations between CA axes 1-3 and PCoA axes 1-3. Only CA2 and PCoA2 were found to be significantly correlated (p = 0.03, rho = -0.27; Spearman correlation); all other comparisons are not significantly correlated. B) Correlations between CA axes 1-3, and Shannon diversity and Faith’s phylogenetic diversity. No significant correlation observed between all comparisons (p > 0.05, Spearman correlation).

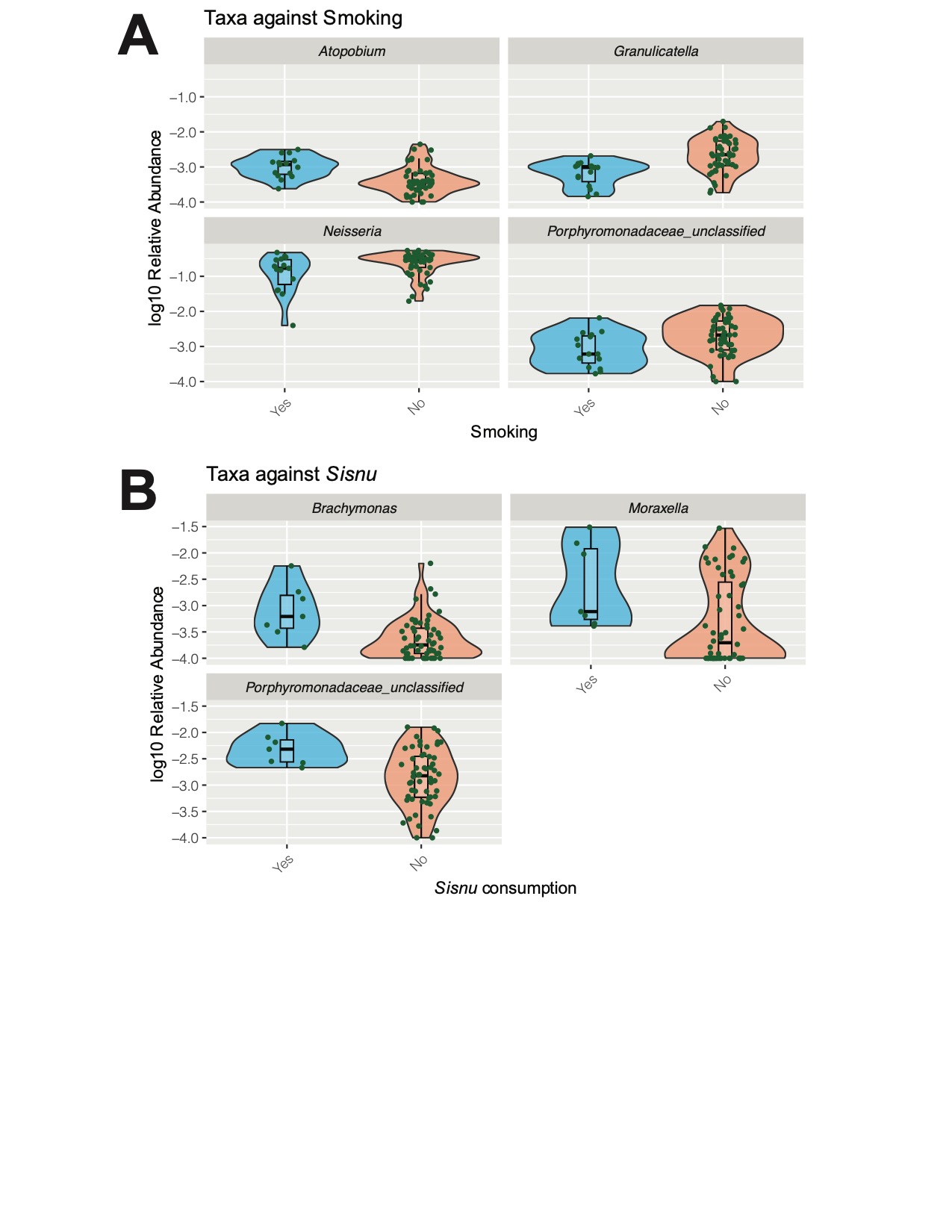

**S11 Figure: Smoking and *sisnu* are associated with several differentially abundant taxa**

A) *Granulicatella*, *Neisseria, Porphyromonadaceae_unclassified*, and *Atopobium* are significantly associated with smoking, prior to multiple test correction (p = 0.006, p = 0.032, p = 0.033, p = 0.023; respectively). Abundances of *Granulicatella*, *Neisseria,* and *Porphyromonadaceae_unclassified* are increased in non-smokers (No), whereas that of *Atopobium* is increased in smokers (Yes). Taxa were log10 transformed for visualization. B) *Brachymonas, Moraxella*, and *Porphyromonadaceae_unclassified* are significantly associated with *sisnu* consumption, prior to multiple test correction (p = 0.014, p = 0.019, p = 0.012; respectively). Abundances of the aforementioned taxa are increased in individuals that consume *sisnu* (Yes).

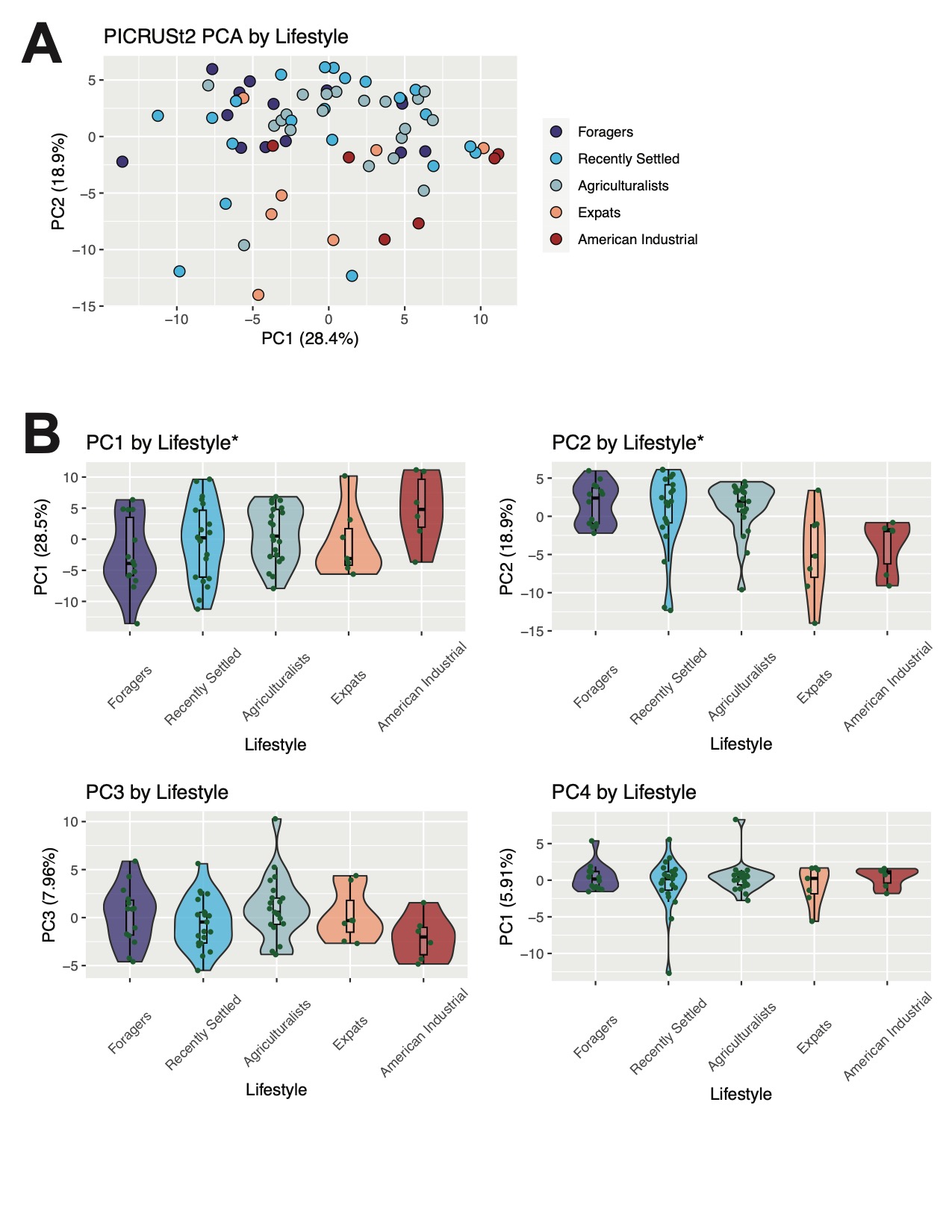

**S12 Figure: Predicted functional abundance significantly differs by lifestyle**

Taxonomic function was predicted via PICRUSt2 from the microbiome data. A) Predicted functional abundance varies by lifestyle (p = 0.0036; PERMANOVA) B) Distribution of individuals along PC axes 1-4, grouped by lifestyle. The distributions of individuals along PC axes 1 and 2 follow the lifestyle gradient (p = 0.049, p = 0.0064, respectively; Jonckheere-Terpstra test) as indicated by *. All other axes do not follow the lifestyle trend (p > 0.05).

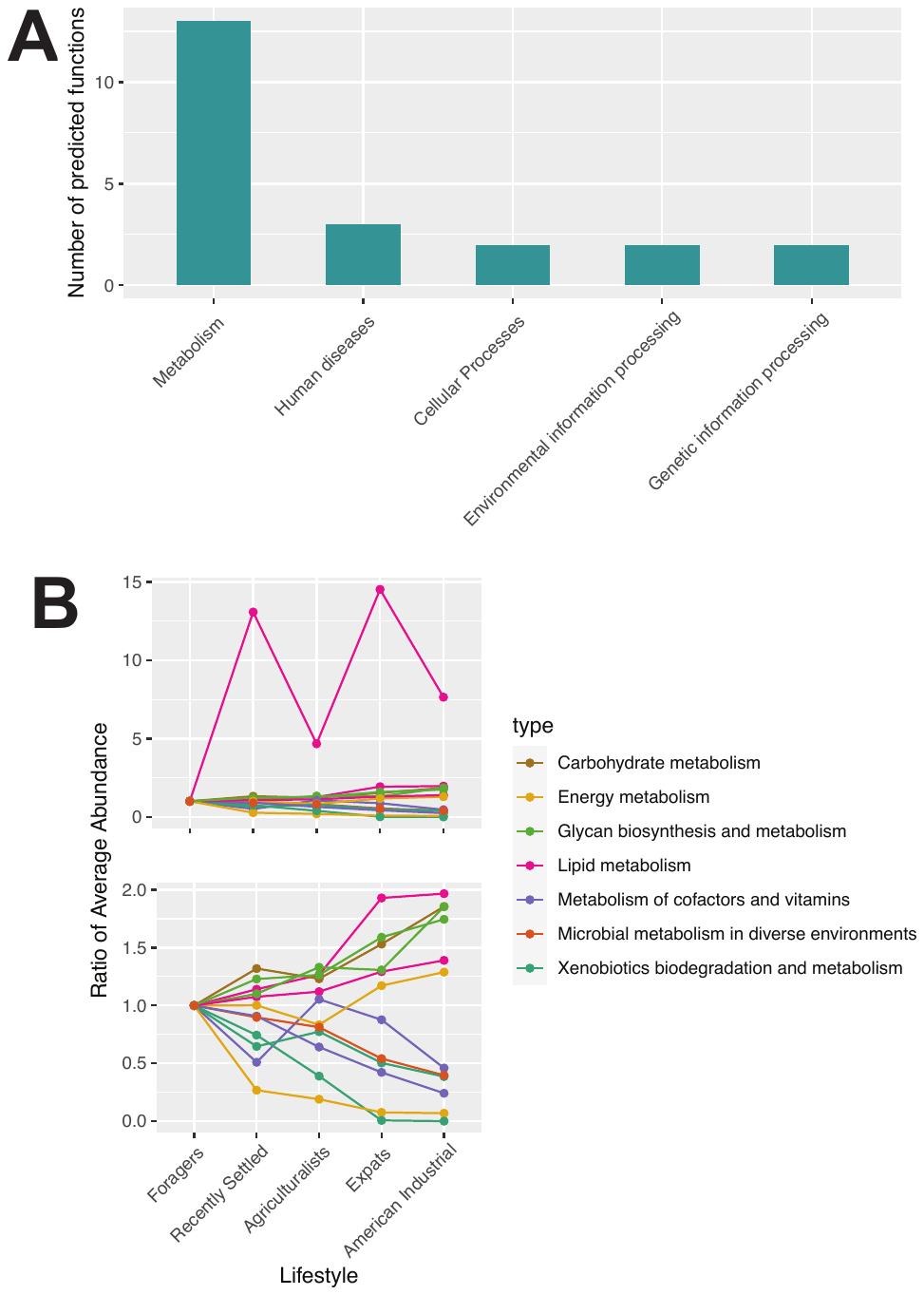

**S13 Figure: Metabolism pathways form the majority of the significantly differentially abundant predicted functions from PICRUSt2**

Metabolism pathways found in the PICRUSt2 predicted functions that are significant prior to multiple test correction. Pathways are classified into five overarching classes and then several smaller subclasses. A) 13/22 of the pathways are classified as the metabolism class. B) Metabolism pathways are categorized into 7 subclasses. Ratios for each pathway were calculated by dividing the average relative abundance within the lifestyle group by the average relative abundance within foragers. (Top) Abundances of lipid metabolism, carbohydrate metabolism, and glycan biosynthesis and metabolism increase with industrialization. Abundances of metabolism of cofactors and vitamins, microbial metabolism in diverse environments, and xenobiotics degradation and metabolism decrease with industrialization. (Bottom) Same pathways, except without Glycerolipid metabolism for visualization purposes.

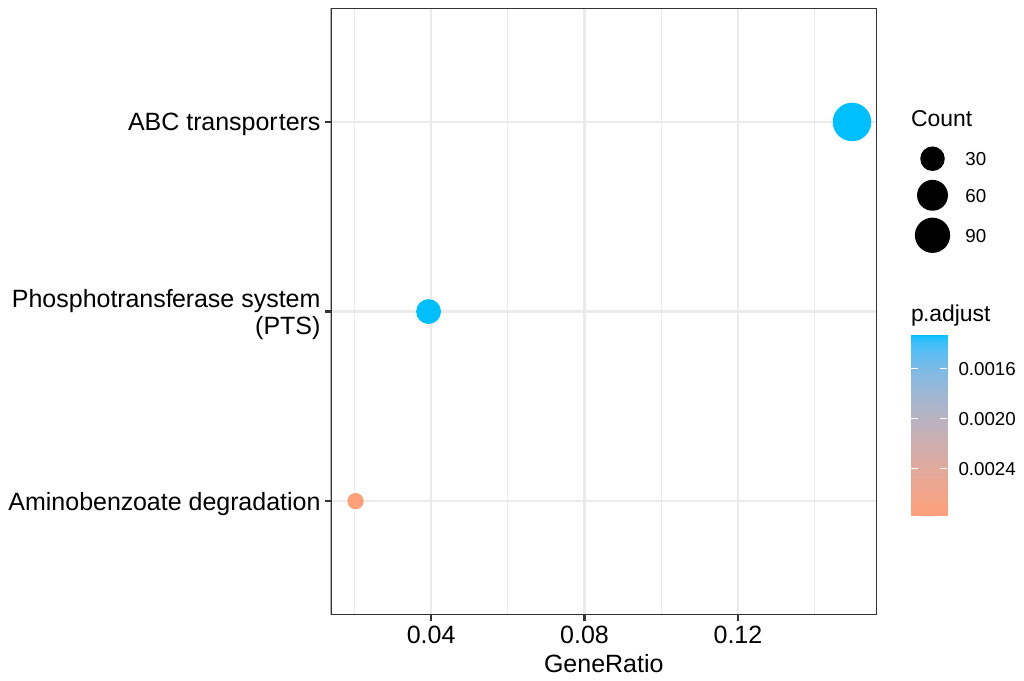

**S14 Figure: Microbiome functional enrichment analysis**

Microbial predicted functions that are enriched within the nominally significant pathways identified from the differential abundance analysis between lifestyles. Pathways with the highest gene ratio (a measure of enrichment) are shown and colored based on significance (p-value corrected with the Benjamini-Hochberg method). Size of the dot indicates unique gene count. In general, transporter proteins and photosynthesis-related pathways are enriched.

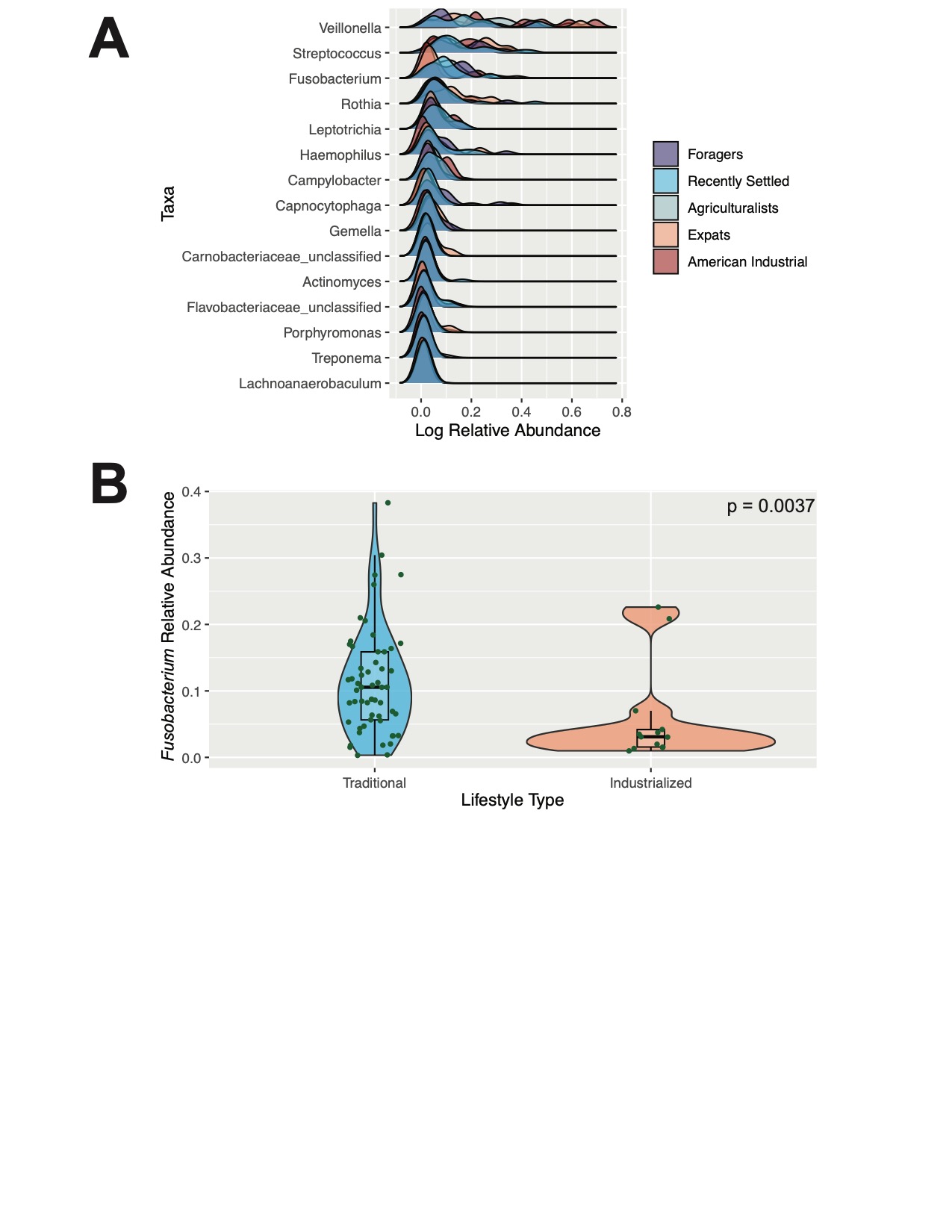

**S15 Figure: *Fusobacterium* contributes to predicted platinum resistance, which varies by lifestyle**

A) Top 15 taxa that most contribute to predicted platinum resistance. Abundance of each taxon (log transformed for visualization) is shown, with individuals grouped by lifestyle. B) Abundance of *Fusobacterium* contributing to platinum resistance significantly differs between the traditional and industrialized populations (p = 0.0037; Kruskal-Wallis test). American industrialists and Expats were categorized as industrialized, whereas the Foragers, Recently Settled, and Agriculturalists are traditional.

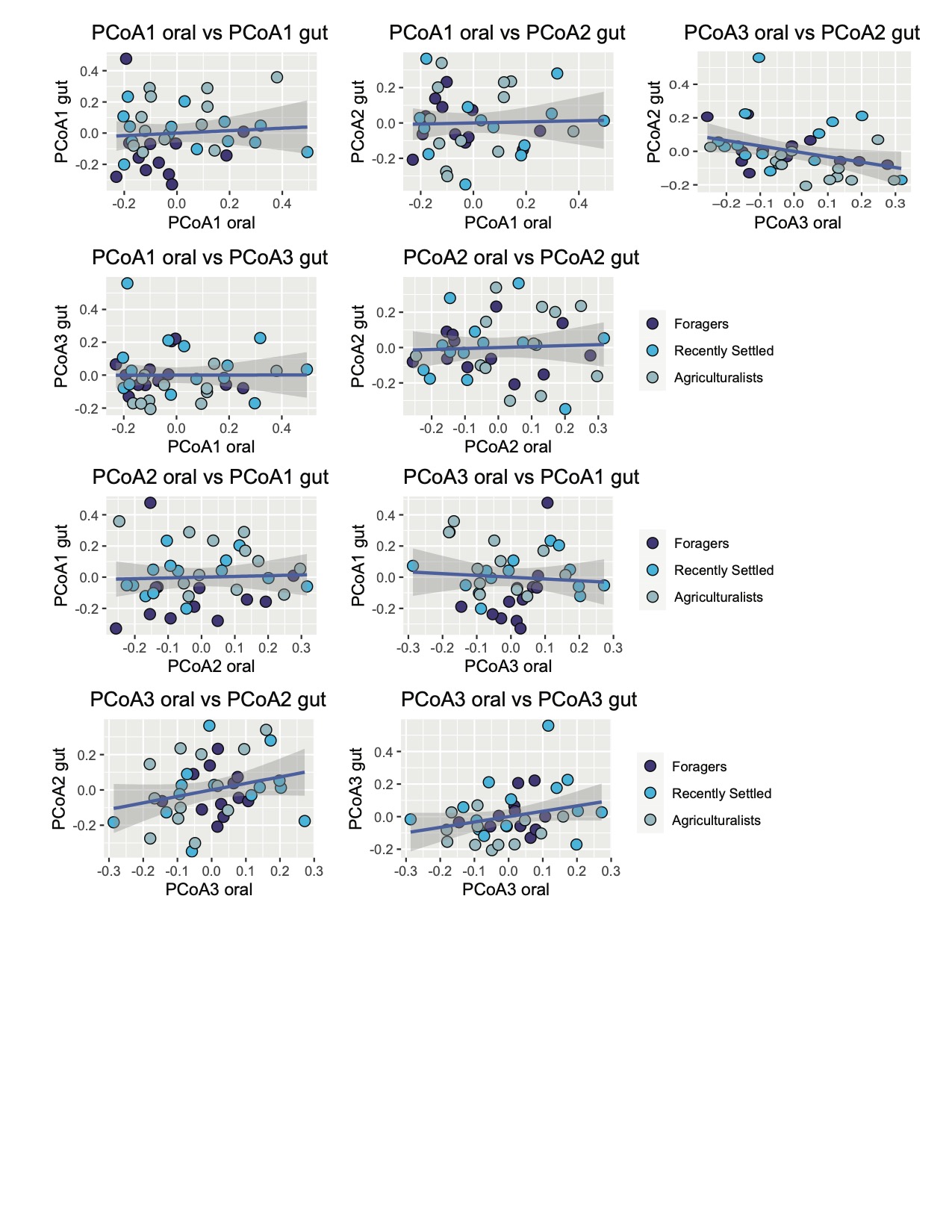

**S16 Figure: Correlations between the top three oral and gut microbiome PCoA axes**

Correlations between the beta diversities of the oral and gut microbiomes, as calculated using the top 3 PCoA axes from each biome (Bray Curtis). Only the relationship PCoA2 from the oral microbiome and PCoA3 from the gut microbiome is significantly correlated (p = 0.013, rho = -0.4; Spearman correlation). All other relationships depicted here are not significant (p > 0.5; Spearman correlation).

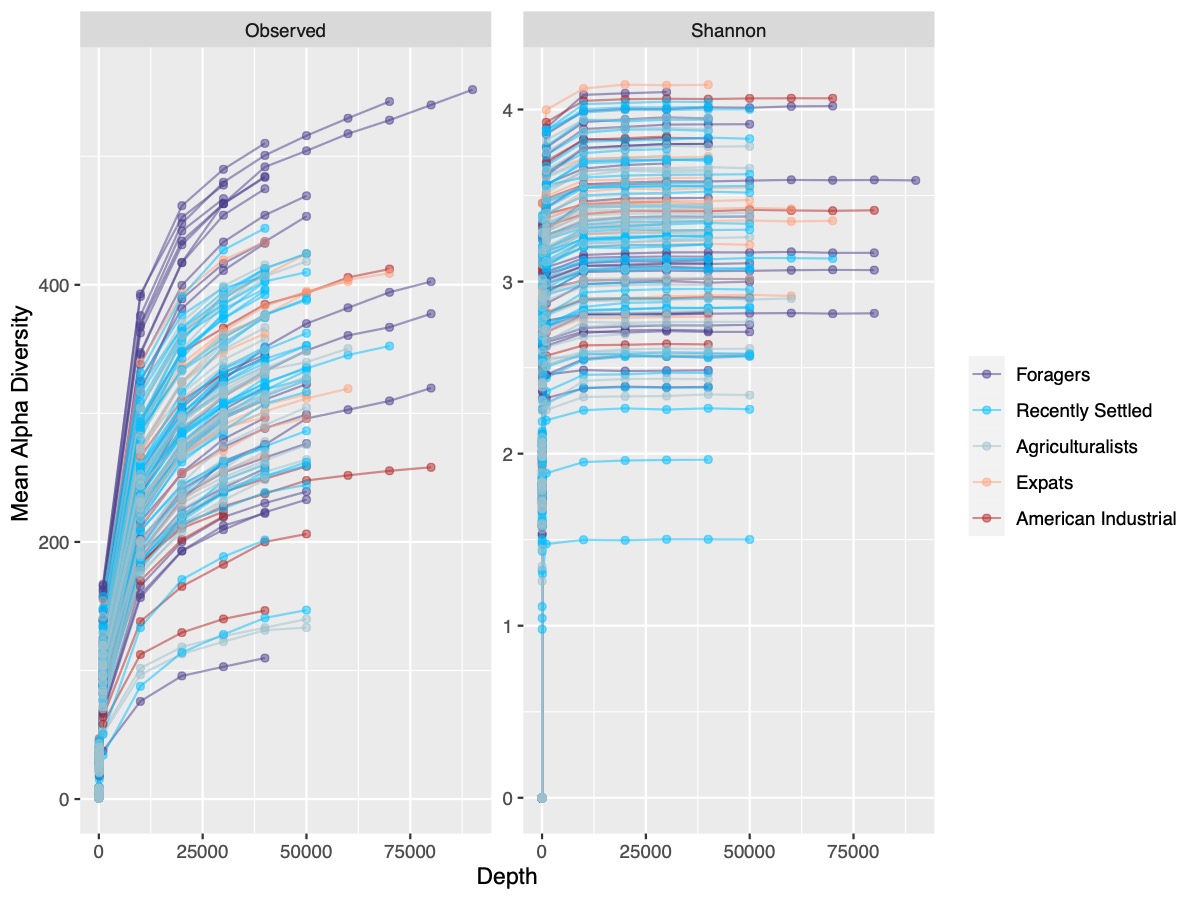

**S17 Figure: Rarefaction curves**

Rarefaction curves showing the mean number of taxa (Observed - left) and Shannon’s diversity (right) with increasing rarefaction depth. Each curve represents an individual and they are colored by lifestyle. Reads were subsampled at a designated rarefaction depth and alpha diversity was calculated from the subsampled reads. This was repeated 10 times and the average was calculated across the 10 trials to account for randomness in rarefaction. This process was repeated at each rarefaction depth, which increased in powers of 10 until 10000, at which point rarefaction depth was calculated at 10000 read increments.

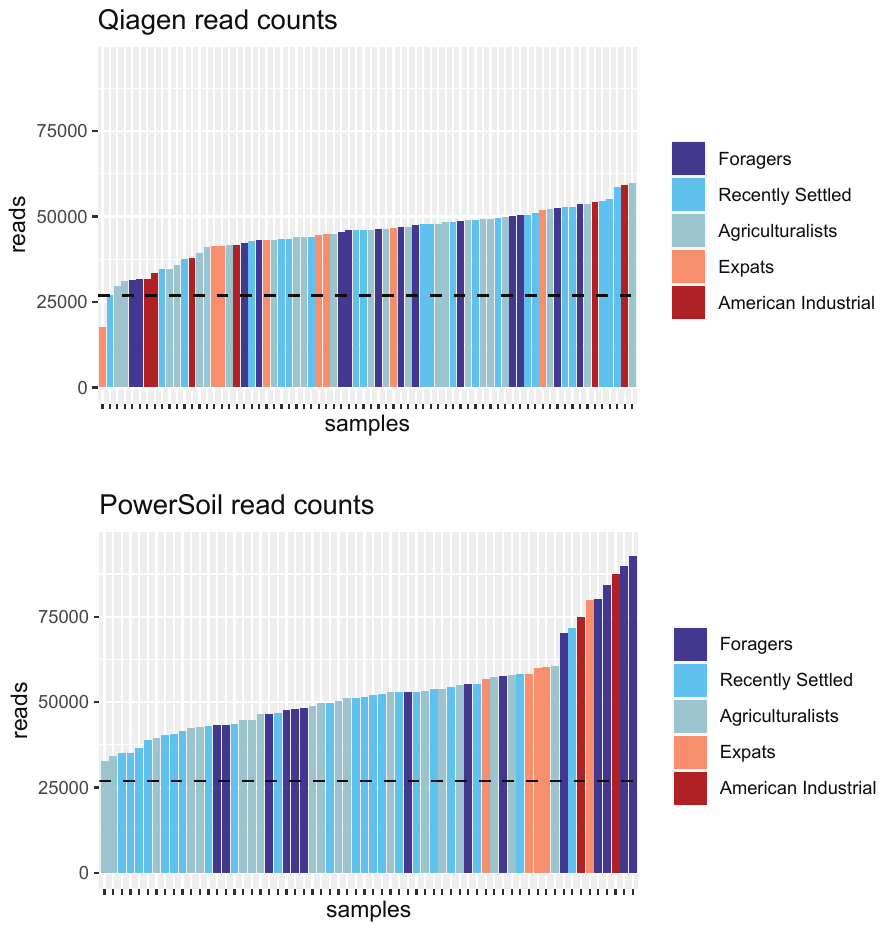

**S18 Figure: Read depth across samples after read QC**

Read depth across all samples after read QC. Samples are ordered from lowest to highest read depth, colored by lifestyle. The black dotted line indicates rarefaction depth.

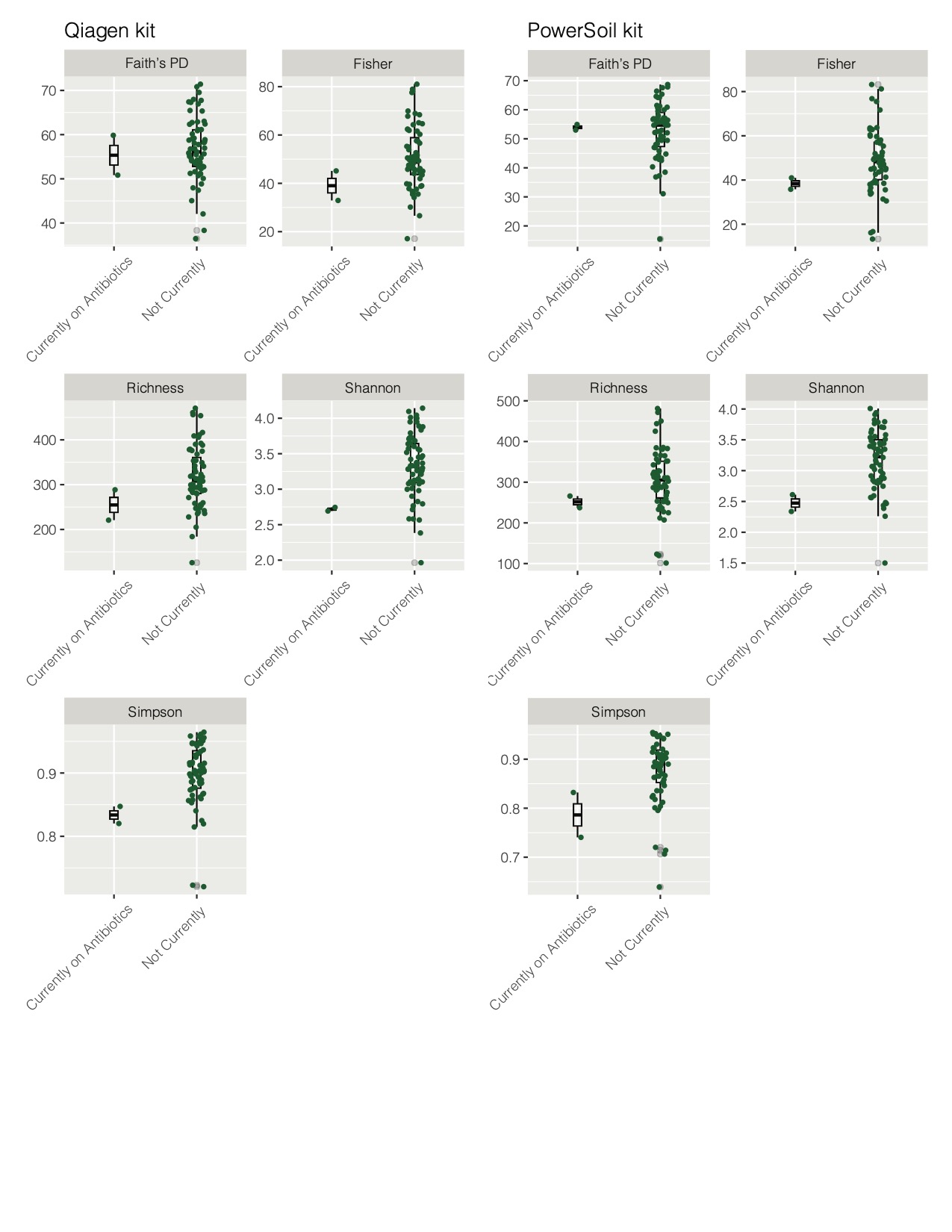

**S19 Figure: Comparison of antibiotic use on oral microbiome diversity**

Current antibiotic use was marginally associated with decreases in oral microbiome diversity as measured by Shannon diversity for samples extracted by the Qiagen kit (p = 0.048, Kruskal-Wallis), but not for other metrics or for samples extracted using the PowerSoil kit (p > 0.05). Due to low sample size of individuals currently on antibiotics, this comparison may be underpowered and prior work suggests that current antibiotic use may have a role in the oral microbiome. Any individuals taking antibiotics were removed for downstream analyses (n = 2).

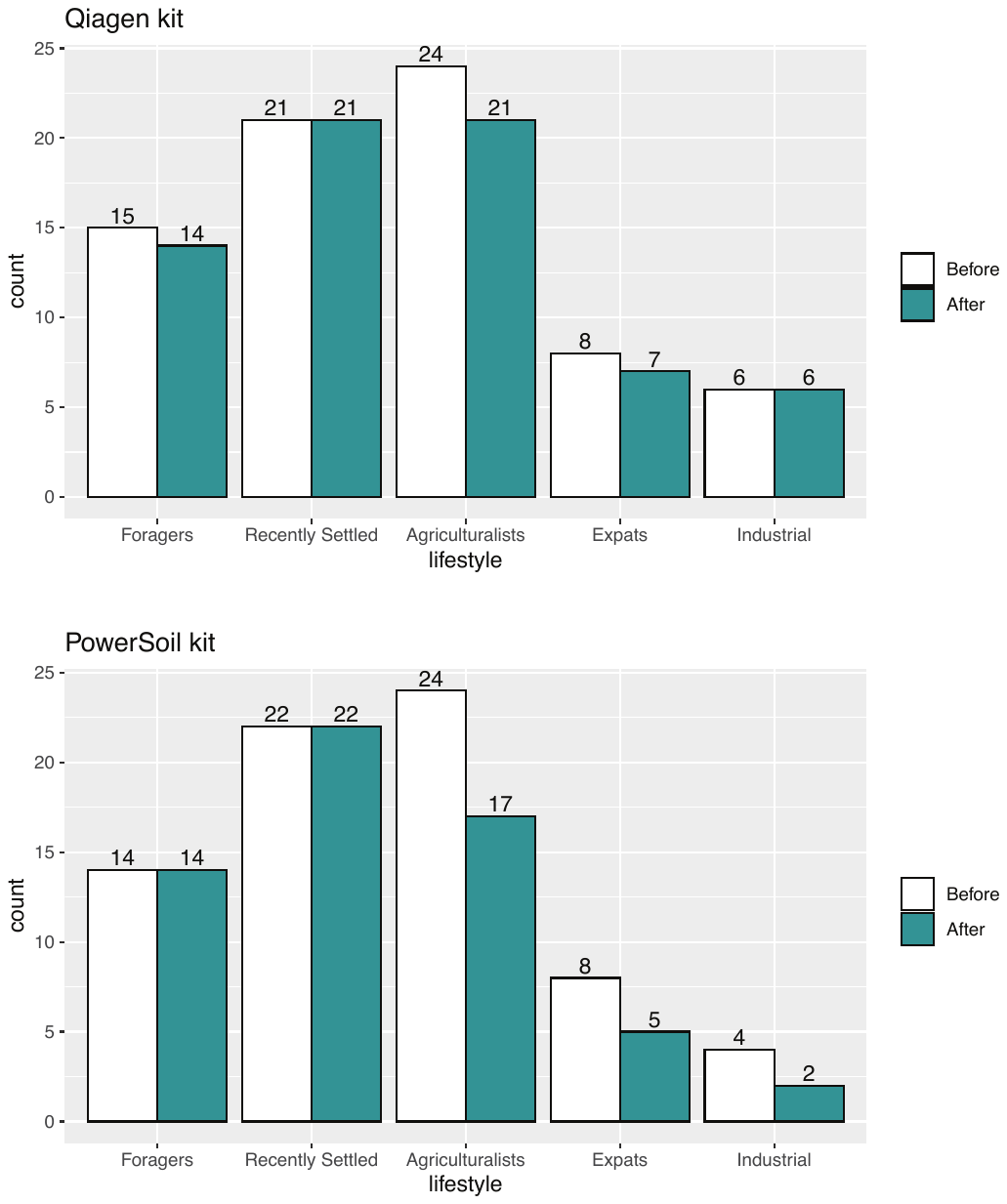

**S20 Figure: Final sample size per lifestyle per extraction kit**

Final sample size per lifestyle per kit before and after quality control preprocessing. Samples are separated by lifestyle and color of bars indicates sample size before and after preprocessing. The post-QC result sample sizes are 69 samples extracted by the Qiagen kit (top) and 60 samples extracted by the PowerSoil kit (bottom).

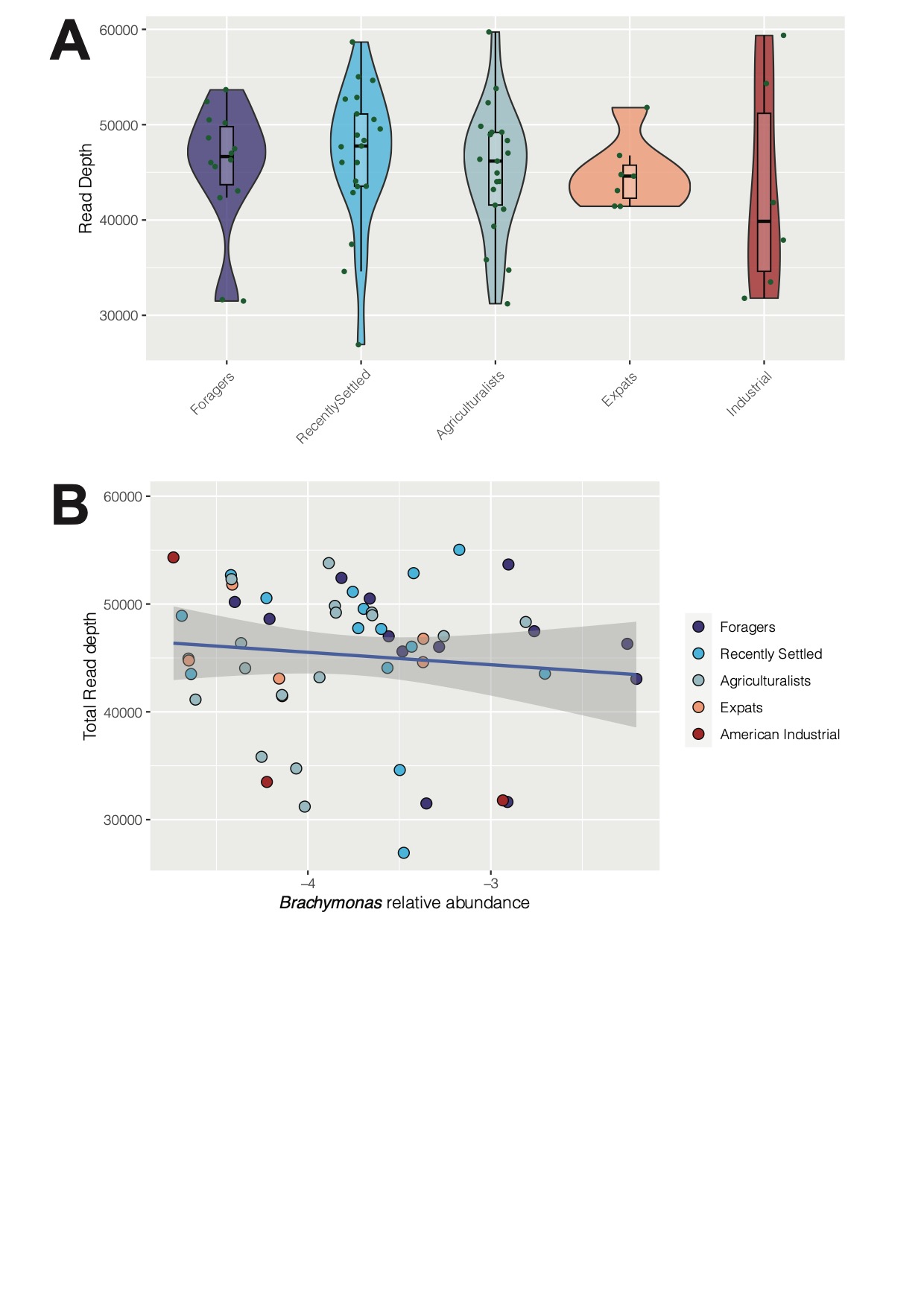

**S21 Figure: Read depth against lifestyle and *Brachymonas* relative abundance**

A) Read depth is not associated with lifestyle (p = 0.63, Kruskal-Wallis). B) *Brachymonas* relative abundance is not correlated with sample read depth (p > 0.05, rho = -0.036, Spearman correlation).
